# Cortical Functional Connectivity in Mouse Models of Early Blindness: Enucleation vs. Anophthalmia

**DOI:** 10.1101/2025.07.17.665343

**Authors:** G. Laliberté, D. Boire

**Author notes:** Corresponding author: Denis Boire.

## Abstract

Early sensory deprivation drives large-scale reconfiguration of cortical networks, yet we still lack a clear understanding of the relative contributions of early visual experience versus spontaneous prenatal retinal waves on the establishment of the cortical network. We compared two mouse models of congenital blindness: neonatal enucleation and congenital anophthalmia, across two genetic strains (C57Bl/6J and ZRDBA) using mesoscopic calcium imaging of spontaneous activity and graph-theoretical analysis. Spectral analyses revealed localized strain-specific increases in infraslow and low delta power following visual deprivation, with C57Bl/6J enucleated and ZRDBA anophthalmic mice exhibiting a more generalized nodal increase. Concomitantly, the functional network organization was redirected toward medial higher-order visual areas, the associative retrosplenial cortex, and somatosensory regions, while the primary and lateral visual cortices exhibited reduced influence and integration within the modular architecture. Notably, ZRDBA groups showed limited global changes to their cortical network. However, anophthalmic ZRDBA mice, lacking prenatal retinal waves, exhibited connectivity patterns more akin to enucleated C57Bl/6J than to their enucleated littermates, highlighting the instructive role of spontaneous prenatal retinal activity. These findings support a connectivity-constrained, experience-dependent model in which preexisting structural pathways guide diffuse, resilient reorganization following sensory loss.

## Introduction

Individuals with visual impairments often display increased sensitivity to intact sensory modalities such as touch (Sadato et al. 1996; Goldreich and Kanics 2006; Ptito et al. 2012), auditions (Lessard et al. 1998; Roder et al. 1999; Poirier et al. 2006; Collignon et al. 2007; Bedny et al. 2011), and olfaction (Renier et al. 2013). These hypersensitivities to the intact sensory modalities in early blind patients coincide with the crossmodal activation of visual cortical areas (Bedny *et al*. 2011; Ptito *et al*. 2012; Renier *et al*. 2013; van der Heijden et al. 2019). Moreover, the functional significance of these crossmodal activation is demonstrated as transcranial magnetic inactivation of these visual areas reduce the crossmodal discriminative capabilities in early blind individuals (Amedi et al. 2004; Collignon et al. 2009).

Together, these findings reflect experience-dependent reconfigurations of brain connectivity. In early blind individuals, resting-state functional connectivity (rsFC) between primary visual cortex (V1) and primary sensory or motor cortices is diminished, while connectivity with frontoparietal cognitive networks is enhanced (Yu et al. 2008; Burton et al. 2014; Heine et al. 2015; Bauer et al. 2017; Abboud and Cohen 2019). These changes suggest a functional redirection of V1 away from sensory integration towards alternative pathways of higher-order processing (Liu et al. 2007; Bedny *et al*. 2011; Watkins et al. 2012; Burton *et al*. 2014; Wang et al. 2014; Striem-Amit et al. 2015; Kanjlia et al. 2016).

Interestingly, similar connectivity profiles are observed in sighted newborns who lack visual experience. This suggests that that long-range cortical functional connectivity is initially sculpted by intrinsic mechanisms and later refined through sensory input (Tian et al. 2023). In contrast to blind adults, sighted adults display stronger functional connectivity between V1 and non-visual sensory cortices, pointing to a dual influence of development and sensory experience in shaping cortical architecture (Tian *et al*. 2023).

Within this framework, the connectivity-constrained experience dependency hypothesis (see Saccone et al. 2024) proposes that preexisting structural constraints shape how deprived cortices can be repurposed following sensory loss. According to this view, in the absence of ascending thalamocortical inputs during development, as in congenital blindness, descending pathways, particularly those from associative cortex, predominate in driving activity within deprived regions (Murphy et al. 2016).

However, the consequences of early sensory deprivation may vary substantially across species, given differences in underlying network architecture. Unlike primates, rodents possess smaller, more fully interconnected cortices, with a reduced hierarchical structure (Harris et al. 2019). They also exhibit direct interconnectivity between modality-specific cortical sensory areas even at early processing stages (Meredith and Lomber 2017; Gamanut et al. 2018; Gamanut and Shimaoka 2022; Magrou et al. 2024). In mice, primary visual cortex receives important descending inputs and displays bidirectional connectivity with associative cortices, suggesting a potential multisensory or integrative roles, even in sighted conditions (Charbonneau et al. 2012; Harris and Mrsic-Flogel 2013; Harris and Shepherd 2015; Petrus et al. 2015; Gamanut *et al*. 2018).

Although dorsal and ventral visual streams have been well established in rodents (Andermann et al. 2011; Wang et al. 2012; Glickfeld et al. 2013; Glickfeld and Olsen 2017; Harris *et al*. 2019; Saleem 2020; D’Souza et al. 2022), these pathways are more tightly interconnected than in primates (Oh et al. 2014; Zingg et al. 2014; Andelin et al. 2018; Gamanut *et al*. 2018; Gamanut and Shimaoka 2022; Magrou *et al*. 2024). This organization aligns with the less modular and more integrated cortical structure of rodents and implies that the impact of visual deprivation may follow differential patterns compared to primates.

In mice, visual cortices respond to non-visual modalities (Oude Lohuis et al. 2024) and are also taken over by the remaining sensory modalities following early blindness (Bonaventure and Karli 1968; Piché et al. 2004; Chabot et al. 2007; Iurilli et al. 2012; Petrus et al. 2014). However, given their dense, nonhierarchical connectivity, it could be expected that rodents exhibit more moderate changes in functional connectivity between sensory-motor areas and between dorsal and ventral visual streams than those reported in primates. The shallow cortical hierarchy and high interconnectivity (Gamanut and Shimaoka 2022) make the mouse brain a valuable model for investigating how early visual deprivation reshapes neural networks, potentially supporting more distributed and resilient reorganization than in the modular primate cortex.

Ultimately, these interspecies differences raise a central question: Does early blindness lead to distinct modes of cortical reorganization depending on species-specific network architecture? The intrinsic interconnectivity of the rodent brain may favor more diffuse and flexible reorganization compared to the more modular and hierarchically structured cortex of primates.

In this context, we used two models of early blindness, neonatal enucleation and congenital anophthalmia, to explore how the timing and presence of early retinal activity shape cortical connectivity. While both models disrupt vision during critical developmental windows (Hensch 2005; Lopez-Bendito et al. 2022), they differ in the preserved early spontaneous retinal activity in the enucleated mice.

The maturation of retinothalamic pathways is initially driven by spontaneous retinal waves (Meister et al. 1991; Wong et al. 1993; Blankenship and Feller 2010; Ackman et al. 2012) which propagate to the thalamus (Mooney et al. 1996; Weliky and Katz 1999) and reach the cortex (Dupont et al. 2006; Hanganu et al. 2006; Colonnese 2014). Prenatal retinal (Cang et al. 2005) and thalamic activity (Anton-Bolanos et al. 2019) are critical for proper formation of cortical maps. Since thalamocortical projections reach the cortical plate around birth in mice (Miller et al. 1993), spontaneous retinal activity can still contribute to early visual structuring in neonatal enucleated mice. In contrast, anophthalmic mice in which the eyecup does not form (Chase and Chase 1941), receive no spontaneous retinal activity, leading to alter thalamocortical and intracortical connectivity during development (Godement et al. 1979; Rhoades et al. 1984, 1985; see also Olavarria et al. 1988; Laramée et al. 2014).

While crossmodal plasticity is well documented across species, the developmental mechanisms driving these changes remain poorly characterized. In particular, the respective contributions of prenatal and early postnatal input to the establishment of long-range cortical wiring are not yet well defined.

In this study, we seek to address these gaps by employing mesoscopic calcium imaging of spontaneous activity in two complementary mouse models of early blindness: neonatal enucleation and congenital anophthalmia. By directly comparing these conditions, we aim to disentangle the influence of spontaneous prenatal sensory-driven activity from that of early postnatal deprivation. Together, these insights highlight the need for controlled experimental models to parse how timing and sensory experience interact with intrinsic network architecture to shape cortical reorganization. In doing so, our work offers new perspectives on connectivity-constrained, experience-dependent plasticity.

## Material and Methods

### Mice

The Animal Care Committee of the University of Quebec at Trois-Rivières (CBSA, protocol DB17) approved all procedures, which adhered to the guidelines of the Canadian Council on Animal Care.

Two mouse strains were used in this study, C57Bl/6J and ZRDBA mice from our animal facility. The ZRDBA mouse line (Touj et al. 2019), derived from crossing sighted DBA-6 with the anophthalmic ZRDCT/An mice (Chase and Chase 1941) which features a key genetic mutation at the *ey1* locus on chromosome 18, crucial for developing structures like the retina (Tucker et al. 2001). Offspring from this cross can either be sighted or anophthalmic, depending on the genetic inheritance of the mutation, allowing for a direct comparison of anophtalmia and early enucleation within a single strain and even littermates.

Monocular deprivation is widely used to study experience-dependant cortical plasticity and has a strong genetic background and heritability (Heimel et al. 2008). The amplitude of ocular dominance shift was reported to be twofold greater in C57Bl/6J compared to the DBA mice (Heimel *et al*. 2008). ZRDBA mice were produced by crossing DBA with ZRDCT/An strain (Chase and Chase 1941; Chase 1942, 1944; Tucker *et al*. 2001) and likely inheriting the more moderate cortical plasticity from DBA mice compared to the C57Bl6J mice. Both C57Bl/6J sighted/enucleated mice and ZRDBA sighted/enucleated/anophthalmic were studied to evaluate the genetic constraints on the cortical plasticity induced by sensory loss.

Twenty-two mice were used in this analysis, distributed into five experimental groups based on strain and eye status. These groups included sighted (n=5) and enucleated (n=3) C57Bl/6J mice, as well as sighted (n=5), enucleated (n=4), and anophthalmic ZRDBA (n=5) mice.

All mice were weaned at 3 weeks, and, except for neonatal enucleation, experimental procedures begins when they reached 6 weeks of age. Data acquisition was conducted between 3 and 6 months of age. Mice were housed in a room with a 12-hour light cycle and provided *ad libitum* access to food and water.

### Enucleation

Upon observing the pup’s birth, the dams received a dose of long-action Buprenorphine SR-LAB (1,0 mg/kg, Chiron Compounding Pharmacy Inc., Guelph, ON). Within 24 hours following birth, bilateral enucleation was performed on half the sighted pups under deep hypothermia induced anesthesia. The palpebral fissure was slit open with a scalpel and the eyeball was gently extracted, sectioning the optic nerve. The ocular orbits were filled with Gelfoam (Spongostan, Johnson & Johnson, New Brunswick, NJ, USA) and the eyelids were sealed with tissue adhesive (Vetbond, 3M, St. Paul, MN, USA). Afterward, the pups were warmed until fully awake and returned to their home cage.

### Intravenous Viral Vector Injection

To target neuronal populations for calcium imaging, viral vectors with the AAV2/PHP.eB capsids were employed to deliver plasmids expressing the GCaMP6s calcium reporter, driven by the human synapsin promoter (AAV-hSyn-GCaMP6s; construct-889-aavphp-eb). The synapsin promoter is commonly used for a broad and stable neuron-specific expression in the cortex (Finneran et al. 2021) while minimizing expression in glial cells (Kugler et al. 2003; Jin et al. 2016). The viral vectors were injected into both C57Bl/6J and ZRDBA mice, aged 6-8 weeks. The viral vectors, provided by the Canadian Neurophotonics Platform Viral Vector Core Facility (RRID:SCR_016477), were delivered via the caudal vein, with a total volume of 300 μL of sterile phosphate-buffered saline (PBS) containing the viral suspension (6.0 x 10^11^ genome copies).

The PhP.eB serotype ability to cross the blood-brain barrier (Chan et al. 2017; Michelson et al. 2019) in both C57Bl/6J and DBA mice (Matsuzaki et al. 2019) allows for delivery into the bloodstream and an efficient neuronal transduction throughout the cortical mantel.

### Implantation of the Chronic Imaging Chamber

At 8-10 weeks of age, all animals were administered a subcutaneous injection of Buprenorphine SR-LAB (1,0 mg/kg, Chiron Compounding Pharmacy Inc., Guelph, ON), anesthetized with isoflurane (induction at 5%, maintenance at 2% in medical O_2_), and secured in a stereotaxic frame. Body temperature was maintained at 37°C using a heating pad and continuously monitored with a rectal thermometer. Additionally, when applicable, the eyes were shielded from corneal dehydration using lubricating veterinary ophthalmic gel (Aventix Pet Eye Lube, Aventix Animal Health, Burlington, ON, Canada). The scalp was shaved and cleaned with chlorhexidine (2% v/v), ethanol (70% v/v), and iodine (16% v/v). Local anesthesia of the scalp was induced with a subcutaneous injection of mixture of Lidocaine 2% and Bupivacaine 0.25% (Lidocaine 7mg/kg, Bupivacaine 3.5mg/kg). The skin covering the skull was then replaced with transparent dental cement (C&B MetaBond, Parkell, Edgewood, NY, USA) with an overlying cover glass (Carolina, Burlington, NC, USA). A titanium bar (Labeo Technologies Inc.; 1.1 g, 25 x 3.2 x 3.2 mm; SKU: 515301) was added for head fixation during recording sessions. Upon surgery completion, mice were given a subcutaneous injection of carprofen (0.5 mg/kg) and individually placed in clean cages for recovery. Subcutaneous injections of carprofen (0.5 mg/kg) were administered at 24-, 48-, and 72-hour post-surgery.

Following a recovery period of 5 days, females were grouped in sets of 3 to 5, while males were paired in cages, separated by clear plexiglass partition to facilitate social interactions while preventing aggression.

### Mesoscale Calcium Imaging Recordings

To minimize potential circadian cycle-related biases, all data acquisition sessions were conducted within the same daily period (between 8 AM and 2 PM). To minimize stress, mice were gradually habituated to head fixation over a period of 5 days: initially for 5 minutes on the apparatus without head fixation, followed by increasing duration of the head fixation (5 minutes, 10 minutes, 20 minutes, and 40 minutes). For the first four days of habituation, brain illumination was off, while on the last day (during the 40-minute head-fixed session), brain illumination was activated. This process effectively reduced signs of stress both within the enclosure and during the experiment, as evidenced by sufficient grooming behaviour, reduced audible vocalization, decreased movements during the head fixation, and stable weight measurements.

During the imaging sessions (Figure 1A), mice were head-fixed in darkness under a wide-field calcium imaging system (OiS200, Labeo Technologies Inc.). Calcium-dependent fluorescence was recorded using a vertically mounted CCD camera (NIKKOR 50 mm f/1.2, Nikon, Minato, Tokyo, Japan) at a spatial resolution of 512 × 512 pixels. Excitation of GCaMP6s was achieved using 472 nm illumination (Cree XLamp XP-E2), the same exposure time of 14.3 ms and a 496 nm long-pass emission filter to minimize non-relevant light contamination.

**Figure 1.**
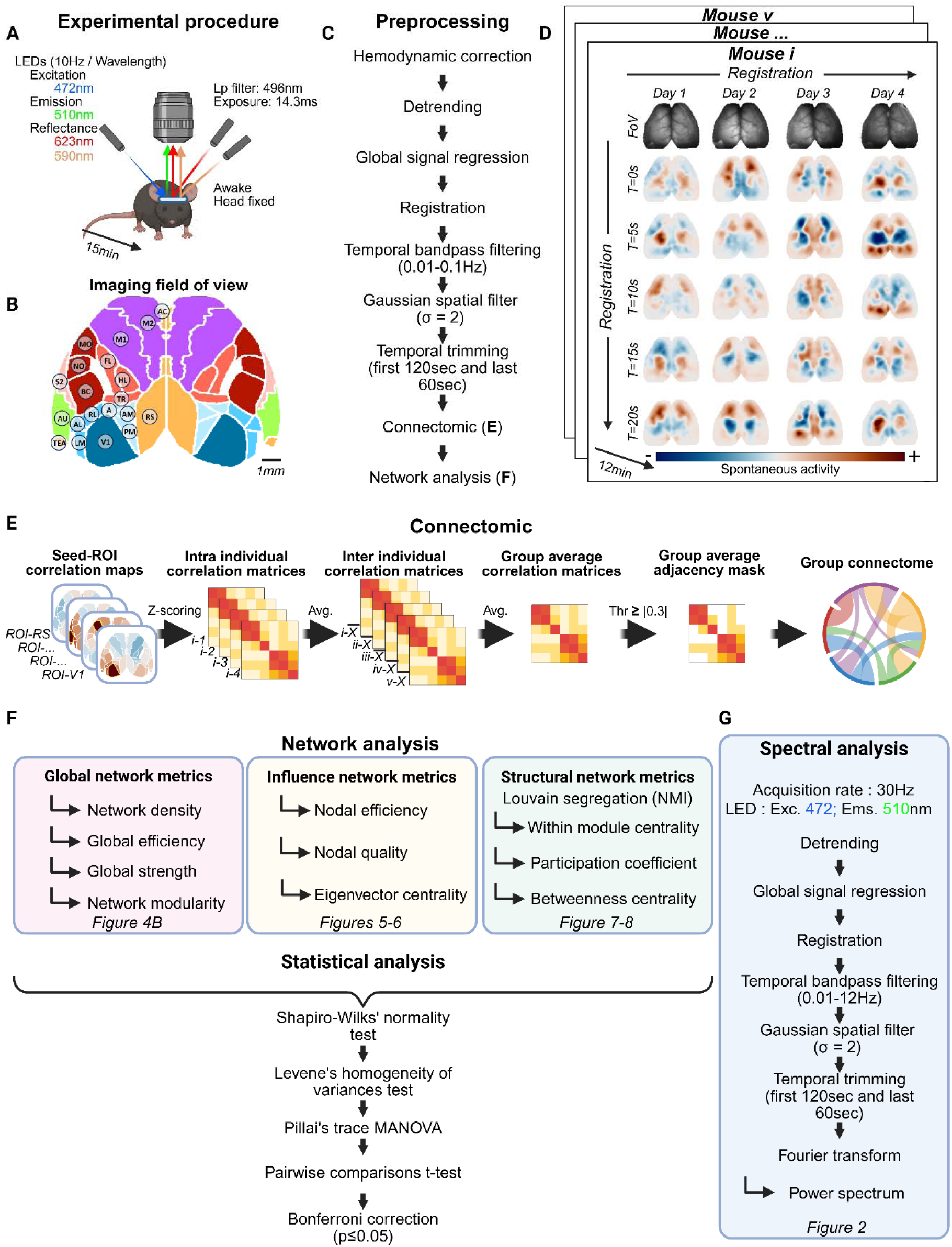
– Summary of the experimental design, preprocessing workflow, and network analysis pipeline for wide field calcium imaging in awake mice. **A.** Schematic of the imaging setup illustrating head fixation, dual-wavelength LED illumination, and the mescoscope used to acquire hemodynamically corrected calcium signals during 15-minute awake recordings. **B**. Cortical parcellation used for region-of-interest (ROI) extraction, based on the Allen Brain Institute Common Coordinate Framework (CCF), depicting all recorded regions within the wide field field of view. **C**. Overview of the preprocessing stream: hemodynamic correction, detrending, global-signal regression, spatial registration across days, temporal band-pass filtering (0.01–0.1 Hz), Gaussian spatial smoothing, and temporal trimming before construction of individual connectivity matrices. **D**. Examples of cross-session spatial alignment for each mouse spontaneous activity epochs followed by a generation of daily correlation maps and group-level averages. **E.** Summary of the connectomic workflow: seed-based correlation mapping, intra- and inter-individual correlation matrices, group average adjacency matrices, and derivation of the group functional connectome. **F.** Network analysis pipeline showing computation of global metrics, nodal influence metrics, and structural community measures obtained from Louvain segmentation, follow-up by MANOVA statistical procedures. **G.** Spectral-analysis stream consisting of detrending, global-signal regression, registration, filtering (0.01–12 Hz), and Fourier transformation to derive ROI-level power spectra (Created in BioRender. LALIBERTÉ, G. (2026) https://BioRender.com/3mfhyg2).

Two acquisition modes were used. For spectral analysis, we recorded resting-state cortex activity for a single 15-minute period acquired at 30 Hz using only 472 nm illumination, without hemodynamic correction. For network construction, each subject underwent four 15-minute sessions in which hemodynamic correction was performed using a cyclic sequence of 472 nm, 590 nm, and 623 nm illumination (Amber and Red LEDs, LZ4-00MA00, OSRAM), enabling extraction of oxy- and deoxyhemoglobin signals (Valley et al. 2020). In this mode, each wavelength was sampled at 10 Hz (30 Hz total sequence rate).

All imaging sessions were conducted in a dark and quiet environment, with the four hemocorrected recordings evenly spaced across a 4–6-week period to ensure stability and reproducibility of the resting-state measurements.

### Data Processing and Analysis of Calcium Signals

Data preprocessing (Figure 1C) was performed with the Universal Mesoscale Imaging Toolbox (UmIT Bipolar-v2.0.0; [an open-source project maintained by LabeoTech; available at <u>labeotech.github.io/Umit</u>]). Images were categorized by wavelength and spatially downsampled to 128×128 pixels (10.7 μm/pixel) using the *run_ImagesClassification* function.

### Spectral Analysis

To determine the specific spectral range in which early blindness alters mesoscale neuronal dynamics and acknowledging that such frequency-specific alterations may partly reflect the characteristics of the imaging approach and calcium reporter, we processed the recordings following the processing pipeline described in Figure 1G. The linear trends were removed using *apply_detrend*, and signals were temporally filtered and normalized with a 0.01–12 Hz band-pass filter (*normalizeLPF*). This range was selected based on methodological constraints: single-wavelength illumination without hemodynamic correction enabled a 30 Hz acquisition rate, yielding a Nyquist limit of 15 Hz; applying a 20% safety margin justified upper cutoffs of 12 Hz, while the 0.01 Hz lower bound removed slow drifts without discarding physiologically relevant fluctuations.

To minimize onset effect, the first 120 seconds and final 60 seconds were removed (*trim_movie*), and spatial noise was reduced via Gaussian filtering (*spatialGaussFilt*, σ = 2). The cortical surface was aligned to the Mouse Allen Brain Atlas top-projection mask (Figure 1B) using *ROImanager*, with dimensions adjusted to the acquisition’s pixel-to-millimeter ratio and each ROI trimmed by 2 pixels to ensure conservative sampling of each ROI. Average time series from 20 right-hemisphere cortical areas (AC, RS, TEA, M1, M2, HL, TR, FL, BC, NO, MO, S2, PM, AM, A, V1, RL, AL, LM and AU) were extracted and subjected to spectral analysis using MATLAB’s *pspectrum* function (R2024b, The MathWorks, Inc., Natick, MA, USA). Group-level spectral power differences were then qualitatively compared between early-blind and sighted cohorts.

### Network Analysis

For the network-construction acquisitions, hemodynamic signal correction was performed following the protocol of Valley *et al*. (2020) using the *run_HemoCorrection* function implemented in UmIT. After detrending (*apply_detrend*), global signal regression (*GSR*) was applied to remove non-neuronal fluctuations, including cardiac and respiratory components (Turley et al. 2017). Each recording was then aligned to a reference frame (*alignFrames*) to ensure intra- and inter-session spatial consistency, with alignment quality verified by an experienced observer.

Temporal band-pass filtering was then applied to isolate target frequency range, specifically 0.01–0.1 Hz for the infraslow frequency (*normalizeLPF*). A Gaussian spatial filter (σ = 2) was used for spatial noise reduction (*spatialGaussFilt*). Acquisitions were trimmed, omitting the initial 120 seconds and final 60 seconds to avoid early habituation and late-session drift (*trim_movie*). Examples of post-processed spontaneous activity waves are shown in Figure 1D.

For connectivity estimation, Pearson correlations were computed between the averaged activity of each of the 20 cortical regions across both hemispheres, using anatomical boundaries defined by the Mouse Allen Brain Atlas top-projection mask (Figure 1B). For every animal, four acquisitions yielded to individual correlation matrix which were transformed into Fisher z-scores (*genCorrelationMatrix*). Each z-score normalized correlation matrices obtained across the hemocorrected sessions were then averaged to produce a single subject-level weighted correlation matrix.

To derive a consistent network structure across animals within the same experimental group, we next computed a group-level mean correlation matrix by averaging the subject-level matrices. This group-average matrix was threshold between 0.1 and 0.4 (step = 0.05) to generate a common adjacency mask (Supplementary Figure 1). The resulting mask was applied back onto each subject’s mean correlation matrix, yielding subject-specific weighted adjacency matrices that were subsequently used for graph-theoretical analyses (Figure 1E).

Graph metrics were computed using the Brain Connectivity Toolbox (Rubinov and Sporns 2010), including density (*density_und*), global efficiency (*efficiency_wei*), strengths (*strengths_und*), eigenvector centrality (*eigenvector_centrality_und*), community structure (*community_louvain*), participation coefficient (*participation_coef*), betweenness centrality (*betweenness_wei*), module degree z-score (*module_degree_zscore*) and normalized mutual information between community partitions (*partition_distance*). All graph visualizations were generated using in-house MATLAB code, except box plots, which were produced using OriginPro 2025b (OriginLab Corporation, Northampton, MA, USA).

### Rationale for Graph-Theoretical Metrics

We characterized how early visual deprivation shapes large-scale cortical organization by selecting a set of graph-theoretical metrics that capture complementary dimensions of network structure. Global metrics, such as network density, global strength, global efficiency, and modularity, summarize the overall pattern of functional coupling and the balance between integration and segregation across the cortical network. These measures are relevant in the context of sensory deprivation, as large-scale coordination and modular differentiation may be selectively altered by the lack of early sensory input. By quantifying the connectivity load (Global strength and Network Density), the efficiency of information transfer (global efficiency), and the extent to which the network retains a modular architecture (Louvain modularity), these global descriptors provide the foundation upon which more localized reorganizations can be interpreted.

To assess how sensory deprivation alter the influence of specific cortical areas within this global framework, we examined for each area nodal quality, efficiency, and eigenvector centrality. These measures provide complementary insights into each region’s contribution to large-scale cortical communication: nodal quality reflects the average weight of a region’s connections, nodal efficiency characterizes its accessibility to the rest of the cortex, and eigenvector centrality identifies regions that hold influential positions within higher-order connectivity patterns. Together, these metrics allow us to determine whether early blindness reduces the hierarchical importance of primary visual cortex, enhances the prominence of associative or somatosensory areas, or distributes influence across high-order visual regions.

Community structure was identified using the Louvain algorithm, which does not require an a priori specification of the number of modules and is therefore well suited for detecting data-driven modular organization in cortical networks undergoing developmental configuration. Because functional reorganization also depends on how cortical areas are embedded within and between modules, we evaluated the structural roles of nodes through within-module degree centrality (WMC), participation coefficient (PC), and betweenness centrality (BeC). Each of these metrics was used to assess structural hubs in the network. The WMC identifies provincial hubs that coordinate processing within a functional community, while the PC highlights connector hubs that link distinct modules. BeC complements these measures by identifying intramodular heavily connected nodes that are also important bridges for intermodular communication.

By integrating global topology, nodal influence, and modular structure, this set of metrics provide a coherent and biologically meaningful framework for capturing the specific large-scale cortical organization induced by early visual loss. Their complementarity enabled the detection of both widespread network-level alterations and more localized shifts in cortical hierarchy or intermodular communication, offering a comprehensive view of how early retinal activity and visual activity shape the functional architecture of the developing cortex.

### Statistical Analysis

Data analysis was performed using IBM SPSS Statistics (Version 29.0.2.0; IBM Corp., Armonk, NY, USA). Intra- and interhemispheric correlation distribution values were computed, and group differences were assessed using Mann–Whitney tests for sighted versus enucleated C57Bl/6J mice and Kruskal–Wallis tests for sighted, enucleated, and anophthalmic ZRDBA mice. Uncorrected Welch’s t-tests were performed on the correlation matrix to explore nodal connectivity alterations in early blind compared to sight mice.

Prior to inferential testing, all graph-derived metrics were evaluated for normality and variance homogeneity using Shapiro–Wilks and Levene’s tests. Most regions of interest satisfied these assumptions within each group, though occasional deviations were present. Because of these rare violations and the modest sample sizes, multivariate analyses based on Pillai’s trace was selected.

Statistical comparisons were performed separately within each strain. For global metrics (density, global strength, global efficiency, and modularity) MANOVA was used to assess the effect of visual status. When the multivariate test was significant, pairwise t-tests with Bonferroni correction (α = 0.05) were applied to identify the specific global properties modified by early blindness.

For nodal metrics, including measures of nodal influence (nodal quality, nodal efficiency, eigenvector centrality) and nodal structural role (WMC, PC, BeC), multivariate analyses examined the combined effect of the visual status and cortical region. Significant interactions were followed by additional univariate analysis across regions to determine which nodes and metrics combination were affected and Bonferroni-corrected pairwise t-tests (α = 0.05) were used to determine which metrics accounted for the observed differences. Nodes with values exceeding one standard deviation above the group mean across the six nodal metrics were classified as structurally or functionally prominent (hubs).

Strain-dependent effects were evaluated by comparing sighted and enucleated animals across the two genetic backgrounds, excluding anophthalmic ZRDBA mice to maintain comparable deprivation conditions. When a multivariate effect was identified, the specific global or regional nodal metrics responsible were determined using the same Bonferroni-corrected pairwise t-test procedure.

## Results

### Spectral analysis

We compared regional power spectra between sighted and blind mice to identify frequency ranges most affected by visual deprivation and to determine whether cortical hubs exhibit preferential spectral alterations (Figure 2). Analyses focused on oscillations in the infraslow (<0.1 Hz), delta (0.1–4 Hz), theta (4–7 Hz), and mu (8–12 Hz) bands, which support large-scale cortical processing and cross-frequency coordination. Moreover, anatomical hub regions play a crucial role in this integration and in driving the synchronization of functional modules (Schmidt et al. 2015).

**Figure 2.**
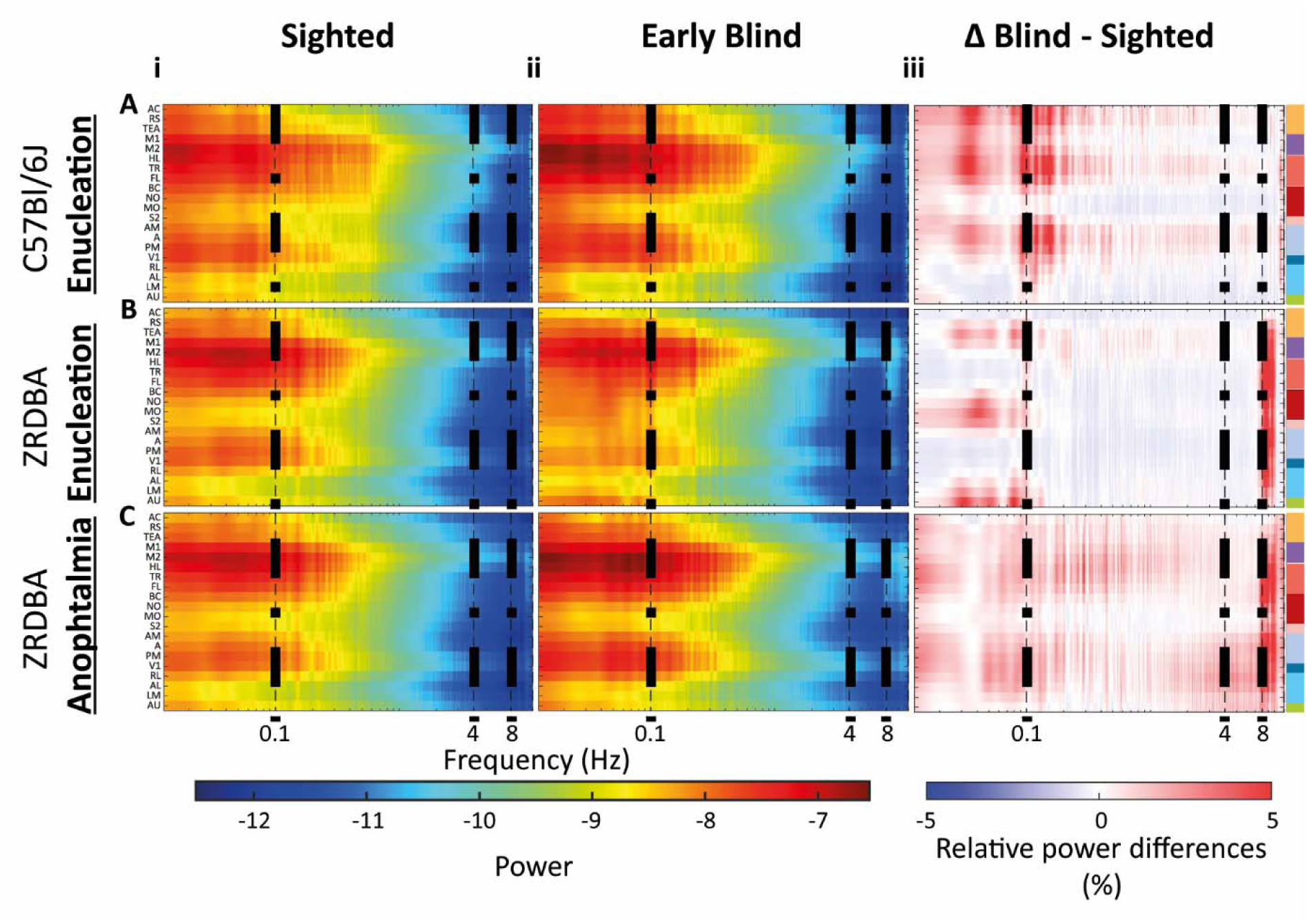
– Resting state Cortical power Spectral Signatures in Sighted and Blind C57Bl6J and ZRDBA Mice Influence of the perinatal enucleation on C57Bl/6J mice (A), on ZRDBA mice (B) and the anophtalmia on ZRDBA mice (C) (i) Smoothed log spectral power in sighted control mice across 20 cortical ROIs on a logarithmic frequency axis. (ii) Corresponding spectral power profiles in early visually deprived mice. (iii) relative power difference maps (Blind – Sighted), highlighting increases (red) or decreases (blue) in power for each ROI. Dashed vertical lines indicate canonical frequency bands: infraslow (<0.1 Hz), delta (0.1–4 Hz), theta (4–8 Hz), and mu (8–12 Hz).

Across groups, spontaneous cortical activity within the recorded range was dominated by infraslow and delta frequencies, with minimal theta and mu contributions (Figure 2). Area-specific spectra across the measured range [0.01–12 Hz] revealed that visual deprivation induced its most pronounced effects in the 0.01–4 Hz frequency range in both strains (Figure 2, columns i–iii). Because the largest and most spatially structured changes occurred in the infraslow band (0.01-0.1 Hz), all subsequent connectivity analyses focused on this range.

In enucleated C57Bl/6J mice compared to sight one, infraslow power increased in associative (AC, RS, TEA), motor (M1, M2), dorsal column–related somatosensory (HL, FL, TR), S2, and medial visual areas (A, AM, PM). Decreases were observed in lateral visual regions (AL, LM), auditory cortex (AU), and somatosensory mouth/nose areas (MO, NO).

In enucleated ZRDBA mice compared to sight controls, infraslow power decreased in AC but increased in RS, TEA, dorsal column–related somatosensory areas (FL, HL, TR), and most visual areas (except AL and LM, which showed increased power). Auditory cortex, trigeminal-related somatosensory areas (BC, MO, NO), and S2 also showed increased power.

While in anophtalmic ZRDBA mice compared to sight controls, mild infraslow power increases occurred in associative (RS, TEA), somatosensory (except MO and NO), and all visual areas, with a slight decrease restricted to AC.

#### Implications for Cortical Integration and Cross-Modal Plasticity

Enucleated C57Bl/6J mice exhibited strong increases in infraslow and delta power across associative, motor, somatosensory, and medial visual hubs, consistent with enhanced large-scale integration. ZRDBA mice displayed more localized modifications, particularly in auditory and trigeminal-related regions. Notably, the infraslow profile of anophthalmic ZRDBA mice more closely resembled that of enucleated C57Bl/6J mice than that of enucleated ZRDBA mice, suggesting the lack of prenatal retinal activity as a stronger effect on the establishment of cortical network frequency profile than the lack of early visual input.

### Functional Connectivity Alterations

We used resting-state spontaneous activity correlations between 20 dorsal cortical areas per hemisphere to generate z-score normalized inter- and intrahemispheric functional connectivity matrices for each experimental group (Supplemental figure 1).

#### Inter- and Intrahemispheric Correlation Distribution

##### C57Bl/6J Enucleated Mice

Analysis of the infraslow connectivity matrix revealed significant alterations of both intrahemispheric (Supplemental figure 2A) and interhemispheric (Supplemental figure 2B) frequency distribution of correlation values. More negative intrahemispheric (Mann-Whitney’s test, p = 0.0028) and positive interhemispheric (Mann-Whitney test, p = 0.001) correlations were observed in enucleated C57Bl/6J mice than in sighted controls.

##### ZRDBA Anophtalmic and Enucleated Mice

In ZRDBA mice, blindness did not significantly affect the frequency distribution of intrahemispheric correlations (Supplemental figure 2C; Kruskal-Wallis test, p = 0.402) in neither enucleated nor anophthalmic group, but the interhemispheric correlation distribution was significantly altered in the blind mice (Supplemental figure 2D; Kruskal-Wallis test, p = 0.009). Corrected multiple comparisons showed a shift towards more negative interhemispheric correlation in the enucleated mice (p = 0.007) than in sighted mice. No significant difference was detected between sighted and anophthalmic mice (p= 0.166) nor between enucleated and anophtalmic mice (p = 0.783).

#### Alteration of the Connectivity Patterns

To explore differences in cortical network dynamics, we initially applied an uncorrected Welch’s t-test (p < 0.05) to compare intra- and interhemispheric resting-state functional connectivity between sighted and early blind mice. This exploratory approach revealed a complex pattern of connectivity alterations, with both increases and decreases across various cortical regions (Figure 3).

**Figure 3.**
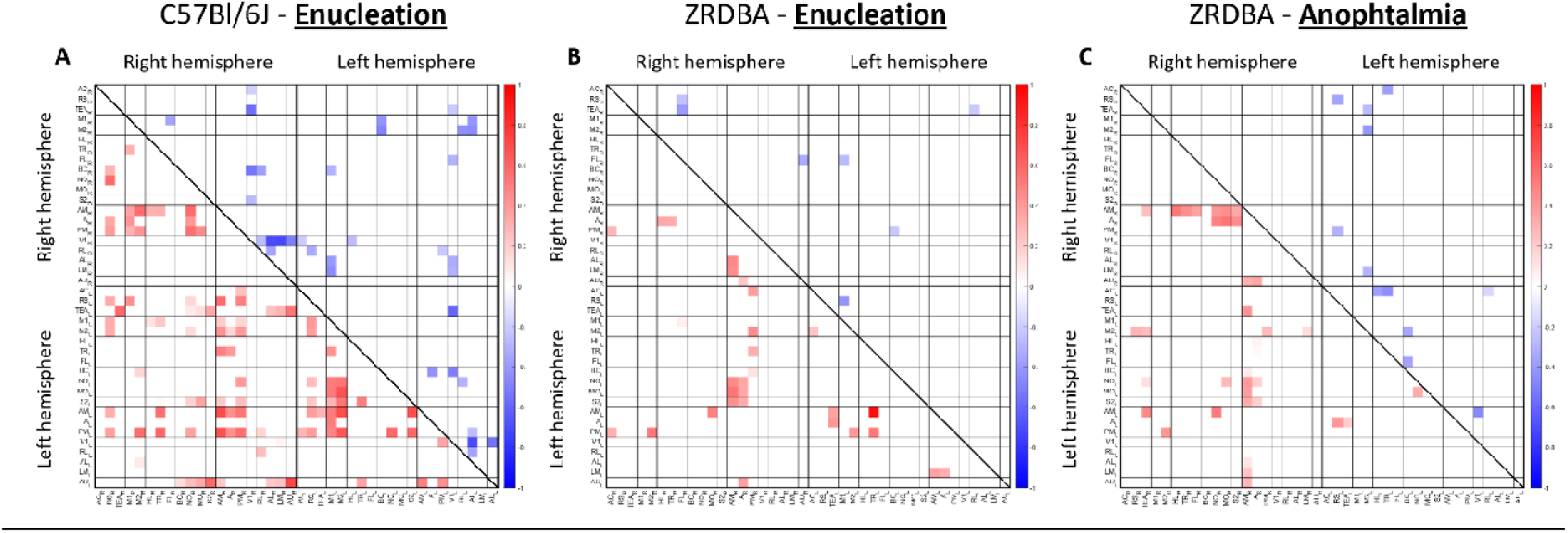
- Altered functional connectivity following early visual deprivation. Significant inter-regional correlation differences (Welch’s t-test, p < 0.05, uncorrected) between early visually deprived mice and sighted controls. Positive differences appear in the lower triangle, negative differences in the upper triangle. (A) C57Bl/6J mice sighted vs. enucleated, (B) ZRDBA mice sighted vs. enucleated and (C) ZRDBA mice sighted vs. anophthalmic. Regions are organized by hemisphere (right and left) and functional domains (associative, motor, somatosensory, visual and auditory).

##### C57Bl/6J Enucleated Mice (Figure 3A)

In C57Bl/6J enucleated mice, homotopic interhemispheric connectivity was increased across associative, visual, and auditory regions, with no observed decreases compared to sighted controls. Heterotopic interhemispheric connectivity also showed widespread enhancements, particularly between medial visual areas and somatosensory, motor, and associative cortices. Additionally, auditory area exhibited stronger interhemispheric connectivity with trigeminal-related and supplemental somatosensory, and lateral visual areas. In contrast, interhemispheric connectivity reductions were confined to motor–sensory interactions, notably between motor and sensory related (BC, AL and LM), and between V1 and lateral visual areas. Intrahemispheric connectivity showed notable increases between medial high-HVAs and associative, motor, and somatosensory cortices. In contrast, decreases were primarily confined to interactions between V1 and lateral visual, somatosensory, and associative regions.

##### ZRDBA Enucleated Mice (Figure 3B)

In ZRDBA enucleated mice, homotopic interhemispheric connectivity was unchanged. Heterotopic interhemispheric alterations were restricted to connections between medial visual areas and trigeminal-related, supplemental somatosensory, associative, auditory, and motor cortices. Intrahemispheric increases were limited between high-order visual areas (HVAS) and associative, auditory and somatosensory areas. Mild decreases were observed between dorsal column-related and associative regions.

##### ZRDBA Anophtalmic Mice (Figure 3C)

In ZRDBA anophthalmic mice, homotopic interhemispheric connectivity decreased in associative and motor cortices. Mild decreases of heterotopic connectivity were observed only between medial HVAs and trigeminal-related, supplemental somatosensory, associative, auditory, and motor areas, as well as between associative and motor regions. Increases of intrahemispheric connectivity were observed between HVAs and associative and somatosensory areas, with no observed reductions.

### Circular Connectomes and Global Network Properties

The impact of neonatal enucleation and anophthalmia on the cortical connectome was assessed by applying a correlation threshold of |r| ≥ 0.3, thereby removing weak or potentially spurious edges and reducing variability driven by noise (Supplementary Figure 3). To quantify large-scale network reorganization, four global graph metrics were extracted: global efficiency, network density, total connectivity strength, and modularity.

Global efficiency captures the average inverse shortest path length, reflecting the network’s capacity for distributed information transfer (Rubinov and Sporns 2010). Density measures the proportion of suprathreshold connections. Total strength represents the sum of all suprathreshold edge weights, indexing the network’s functional robustness (van den Heuvel and Sporns 2011). Finally, Louvain modularity quantifies the extent of community structure, with higher values indicating stronger functional segregation.

Because these metrics capture distinct yet non-redundant dimensions of global network organization, integration (efficiency), connectivity load (density and strength), and segregation (modularity), their combined use within a multivariate framework enables the detection of coordinated, system-level shifts in cortical architecture that may not be apparent when metrics are analyzed separately.

#### Intrahemispheric Network Alterations

##### C57Bl/6J Enucleated Mice Connectome (Figure 4A-i)

Compared to sighted, enucleated C57Bl/6J mice’s V1 lost connections with the somatosensory and auditory cortices. However, V1 established new connections with associative (RS) and motor (M1) V1 maintained its connectivity with HVAs. Enucleation also resulted in the loss of connections between lateral HVAs (AL, LM, and RL) and the medial HVA, PM. Conversely, connectivity increased between other medial HVAs (A and AM) and lateral HVAs (AL and LM).

**Figure 4.**
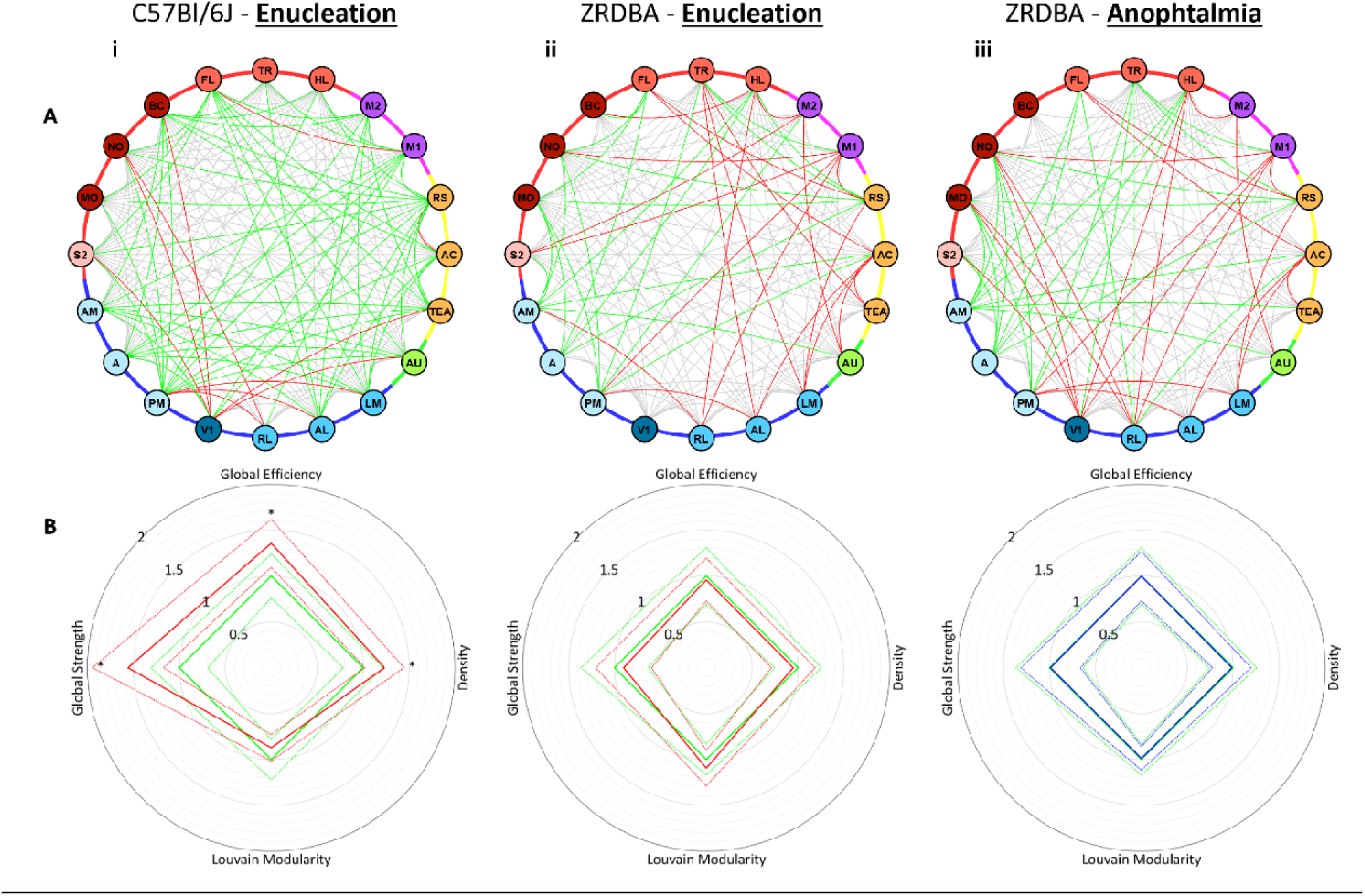
– Differential Circular Connectome Diagram and Global Network Metrics. Influence of the perinatal enucleation on C57Bl/6J mice (**i**), on ZRDBA mice (**ii**) and anophtalmia on ZRDBA mice (**iii**). **A.** The circular connectome diagram of the connectivity patterns between cortical regions (nodes), Suprathreshold z-score (>0.3) connectivity between nodes is depicted with colored edges. Grey links represent connectivity present in both sighted and blind mice, green links indicate suprathreshold connections unique to the blind mice, and red links show connections lost in the blind mice. Perimeter colors represent functional modules as described in Figure 1 (Associative areas (AC, RS and TEA) are yellow. Motor areas (M1 and M2) are in purple, and somatosensory areas in shades of red. Trigeminal sensory regions (MO, NO, and BC) are in dark red, while dorsal column-related regions (FL, HL, and TR) are shown in salmon, with S2 in pink. Auditory area (AU) is represented in green. Visual areas are in shades of blue: V1 in dark blue, lateral visual areas (AL, LM, and RL) in ocean blue, and medial visual areas (A, AM, and PM) in meltwater blue). **B.** Comparison of global network metrics: Four global network metrics—global efficiency (North), network density (East), Louvain modularity (South), and global strength (West)—are compared between sighted (green) and blind groups (red: enucleated, blue: anophthalmic). Metrics are normalized to the mean values of the sighted group (centered at 1). Darker lines represent group means, and lighter lines indicate SEM. Statistically significant differences (p ≤ 0.05, Bonferroni-corrected pairwise tests) are denoted by asterisks (*).

Both lateral and medial HVAs exhibited an overall increase in suprathreshold connections, forming stronger links with other functional domains, including associative, auditory, motor, and somatosensory areas. More specifically, medial HVAs predominantly gained sensory connections from auditory and trigeminal-related somatosensory areas, whereas lateral HVAs primarily integrated sensory inputs from dorsal column-related somatosensory areas. Finally, associative areas, particularly RS, increased its connectivity with all functional domains (associative, auditory, somatosensory, motor and visual).

##### Global Network Metrics Alterations in Early Blind C57Bl/6J Mice (Figure 4B-i)

A Pillai’s trace multivariate test on global network metrics revealed significant differences based on visual status in C57Bl/6J mice (V = 0.872, F [4,4] = 6.828, p = 0.045). Post hoc Bonferroni corrected pairwise comparisons indicated significant increases in network density (p = 0.050), global efficiency (p = 0.015), and global strength (p = 0.007) in enucleated mice compared to sighted controls in the infraslow frequency range.

##### ZRDBA Enucleated Mice connectome (Figure 4A-ii)

In ZRDBA mice, enucleation impacted the cortical network structure differently than in C57Bl/6J mice. Indeed, enucleation did not alter V1 connectivity. However, lateral HVAs lost connections with associative, dorsal column-related somatosensory, and motor areas. In contrast, medial HVAs increased connectivity with trigeminal and dorsal column-related somatosensory areas, as well as with associative regions (AC and RS).

Other changes in connectivity were also observed in associative regions, AC lost connections with auditory areas, whereas RS strengthened its connections with dorsal column-related somatosensory areas.

##### ZRDBA Anophtalmic Mice Connectome (Figure 4-iii)

In anophthalmic ZRDBA mice compared to sighted controls, V1 exhibited a reduction in connections with some somatosensory areas (S2, MO, NO and HL), but contrary to what’s was observed in enucleated C57Bl/6J mice, it retained its connectivity with other sensory areas such as the barrel field cortex (BC) and auditory areas (AU).

Regarding medial HVAs, only PM showed connectivity losses, specifically with the anterior cingulate cortex (AC), the lateral HVAs (AL and LM), and motor area. However, concerning the medial HVAs, A, gained only a connection with RS, whereas AM, established several new connections with associative areas (AC and TEA), dorsal column-related (FL and HL) and trigeminal (MO and NO) somatosensory and auditory (AU) areas. Additionally, connectivity between lateral HVA (RL) and dorsal column-related somatosensory (FL and TR) and associative (AC) areas were enhanced. In contrast, the other lateral HVAs (AL and LM), demonstrated a decreased connectivity with associative areas (AC and RS).

Associative cortices showed other alterations. Notably, AC and RS lost connections with forelimb-related somatosensory areas (FL) and RS gained connections with somatosensory areas (FL, MO and NO).

##### Global Network Metrics Alterations in Early Blind ZRDBA Mice (Figure 4B-ii and 4B-iii)

No significant multivariate effect of visual status was observed in ZRDBA mice (V = 0.385, F [8,18] = 0.536, p = 0.814) for global network metrics. Additionally, Bonferroni-corrected pairwise comparisons revealed no significant differences when comparing sighted to enucleated mice, sighted to anophthalmic mice, or enucleated to anophthalmic mice.

### Nodal Dynamics in Cortical Network Reorganization

To evaluate cortical network reorganization following early visual deprivation, we examined node-level changes using three graph-theoretical metrics: nodal efficiency, connectivity quality, and eigenvector centrality. These metrics respectively capture each region’s capacity for global communication, local connection strength, and integration within highly connected substructures (Lohmann et al. 2010; Rubinov and Sporns 2010). Together, they provide a multidimensional profile of each node’s functional prominence.

#### Reallocation of Cortical Nodal Influences Following Neonatal Enucleation in C57Bl/6J Mice (Figure 5)

Multivariate analysis (Pillai’s trace: V = 0.851, F [57, 420] = 2.920, p < 0.001) confirmed that neonatal enucleation induces widespread reorganization of cortical influence in C57Bl/6J.

**Figure 5.**
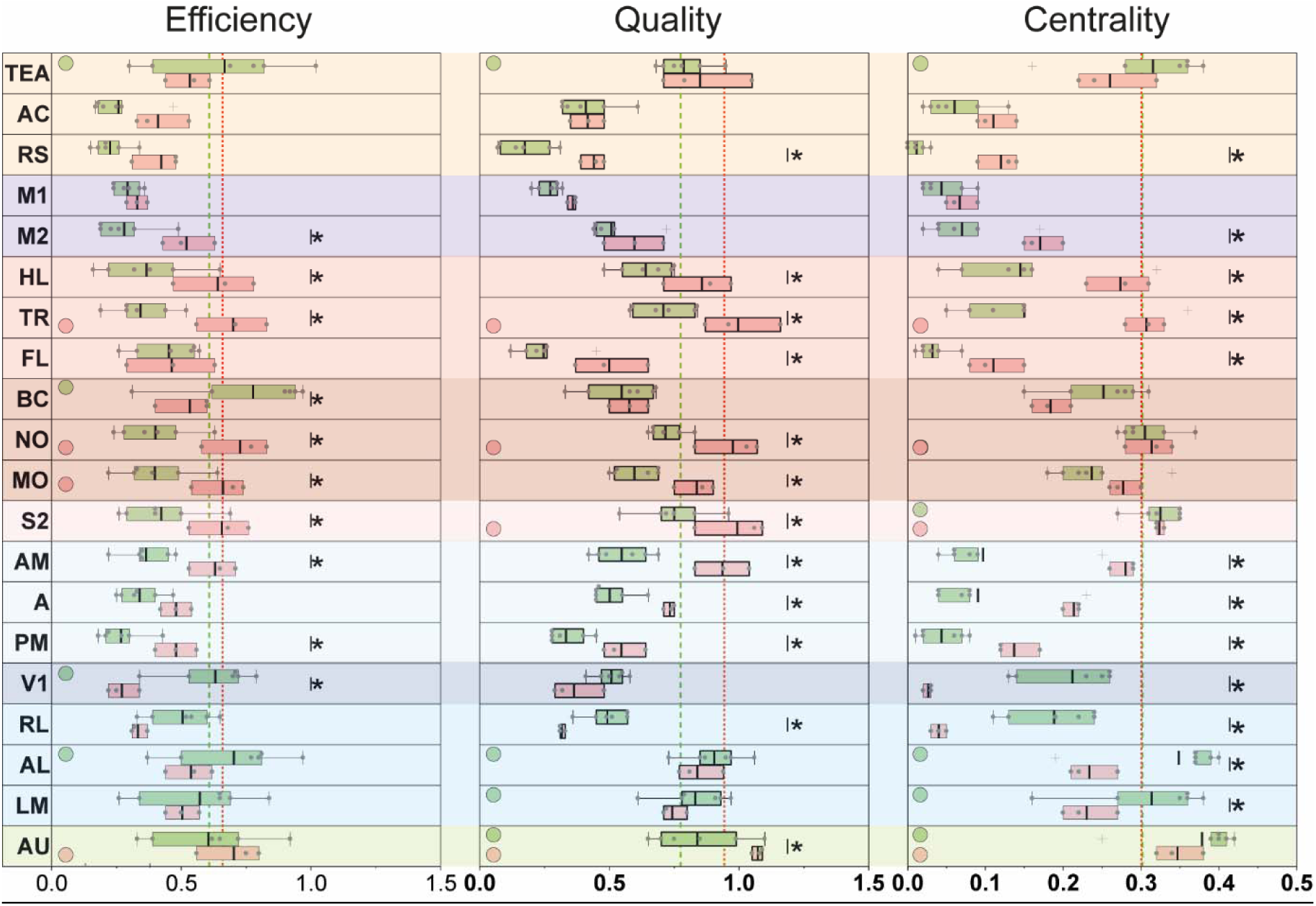
– Effects of Early Visual Deprivation on Cortical Nodal Influence Metrics in C57Bl/6J Mice. Cortical distributions of nodal influence metrics—namely nodal efficiency, connectivity strength, and eigenvector centrality—are shown for sighted (green) and enucleated (red) C57Bl/6J mice. Colored markers denote cortical hubs for each metric, defined as nodes with values exceeding one standard deviation above the network mean (threshold shown as dashed lines). Boxplots display the interquartile range (25th–75th percentiles), with whiskers corresponding to the 95% confidence interval; outliers are represented by crosses and were retained in the statistical analyses, as no methodological or biological justification supported their exclusion. The background color indicates the functional affiliation of each cortical area, following the functional grouping scheme presented in Figure 3. Statistically significant differences between groups, based on Bonferroni-corrected tests (p ≤ 0.05), are marked by asterisks (*).

Nodes with scores exceeding one standard deviation above the group mean on any individual metric were classified as high-impact hubs. Based on their respective metrics, these hubs were categorized as efficient hubs (nodal efficiency), high-quality hubs (connectivity weight normalized by degree), and core hubs (eigenvector centrality). Among visual cortices of sighted C57Bl/6J mice network, AL and V1 were labelled as efficient hubs, while AL and LM were considered as high-quality and core hubs. In enucleated network, all these areas demonstrated lost of their influential hubs’ qualification. Lateral (AL, LM, RL) and primary (V1) visual regions demonstrated the most prominent lost, with V1 exhibiting significant reductions in both efficiency (p < 0.001) and centrality (p < 0.001), while AL exhibited a decline in centrality (p = 0.004). RL also lost influence, with significant decreases in centrality (p < 0.001) and connectivity quality (p = 0.020). In contrast, medial HVAs (A, AM, PM) became more prominent. A showed increased quality (p = 0.002) and centrality (p = 0.002), while AM and PM exhibited significant gains across all three metrics (AM: quality (p < 0.001), centrality (p < 0.001), and efficiency (p = 0.012), PM: quality (p = 0.005), centrality (p = 0.019), and efficiency (p = 0.044)).

In sighted C57BL/6J mice, the barrel cortex (BC) was the only somatosensory region classified as an efficiency hub, whereas the supplemental somatosensory area (S2) qualified as a core hub. Notably, the nose region (NO) retained its designation as a core hub across both visual conditions. In enucleated C57Bl/6J mice, BC lost its efficiency hub status, while multiple somatosensory regions exhibited marked increases in network prominence (MO, S2, TR). The trunk (TR) region emerged as a key driver of network influence post-enucleation, showing robust increases across all three metrics (p < 0.001) and earning classification as efficient, high-quality and core hub. MO gained status as an efficiency hub through elevated nodal efficiency (p = 0.013) and connectivity quality (p = 0.002), while NO showed significant gains in efficiency (p = 0.002) and quality (p < 0.001), qualifying as both an efficient and high-quality hub. S2 maintained its role as a core hub and further gained designation as an efficient and high-quality hub, supported by increased efficiency (p = 0.026) and connection quality (p = 0.001). Hind limb (HL) regions exhibited consistent enhancements across all metrics, efficiency (p = 0.001), quality (p = 0.004), and centrality (p = 0.001), while the forelimb (FL) demonstrated increased quality (p < 0.001) and centrality (p = 0.048). The secondary motor cortex (M2) also gained influence in the enucleated mice cortical network, with significant elevations in efficiency (p = 0.022) and centrality (p = 0.012).

Of the associative and auditory cortices of sighted mice, the temporal association area (TEA) was classified as an efficient, high-quality, and core hub, indicating prominent network integration. However, in the enucleated mice, non-significant decreases across all metrics led to the loss of these classifications. In contrast, the primary auditory cortex (AU) maintained its status as a high-quality and core hub across both visual conditions. While the increase in nodal efficiency in enucleated C57Bl/6J mice did not reach significance, AU nonetheless gained designation as an efficient hub. Notably, AU exhibited a significant increase in connection quality in enucleated mice (p = 0.002). Furthermore, the retrosplenial cortex (RS) demonstrated strengthened network influence in enucleated mice, with significant increase in connection quality (p < 0.001) and centrality (p = 0.007).

To summarize these observations, cross-metric ranking allowed for the descriptive identification of both the most and least influential nodes within each experimental group, as well as those demonstrating shifts in network influence following sensory deprivation. Nodes ranking highest across the combination of all three metrics were designated as influential hubs, representing regions with the greatest overall network impact. In sighted C57BL/6J mice, cortical network influence was predominantly driven by influential hubs such as auditory (AU), associative (TEA), and lateral visual areas (AL, LM), whereas associative (AC, RS), motor (M1), and medial visual (PM) regions exhibited the lowest influence. Following enucleation, the auditory cortex (AU) remained a key driver, accompanied by increased contributions from somatosensory regions (NO, TR, S2). In contrast, associative (AC) and motor (M1) areas persisted among the least influential nodes, now joined by visual regions (V1, AL). Overall, somatosensory (NO, TR) and medial visual areas (AM, PM) demonstrated the greatest increase in nodal influence after visual deprivation, while associative (TEA), lateral visual (AL, RL, LM), and primary visual (V1) areas showed the most pronounced decline.

#### Differential Reorganization of Cortical Nodal Influence in Anophthalmic and Enucleated ZRDBA Mice (Figure 6)

In the sighted ZRDBA mice, cortical network influence was primarily driven by auditory (AU), somatosensory (NO, S2), and lateral visual (AL) regions (influential hubs), while associative (RS), motor (M1, M2), and somatosensory (FL) areas exhibited the lowest influence. In the enucleated ZRDBA mice, influential hubs were composed of auditory (AU) and somatosensory (BC, S2) areas, while associative (AC), motor (M1, M2) and somatosensory (FL) areas were among the least influential. In the anophthalmic ZRDBA mice, auditory (AU) cortex also retained its influential hub rank, accompanied by associative (TEA) and somatosensory (NO, S2) areas, whereas associative (RS) and somatomotor areas (FL, M1, and M2) ranked among the least influential. Cortical network comparison between sighted and enucleated ZRDBA mice revealed pronounced increases in nodal influence ranks for associative (RS) and visual (A, AM, RL) areas, and rank declines of associative (AC, TEA) and somatomotor areas (M1, M2, HL, MO). While comparing sighted and anophtalmic ZRDBA mice, associative (RS), medial visual (A, AM, PM) and somatosensory (TR) areas demonstrated the strongest influential rank gains, while associative (AC), motor (M2) and visual (RL, V1) areas showed the strongest influential rank reductions. Overall, early visual deprivation in ZRDBA mice induced significant although moderate cortical reorganization relative to C57BL/6J mice (Pillai’s trace: V = 0.716, F [114, 660] = 1.814, p < 0.001).

**Figure 6.**
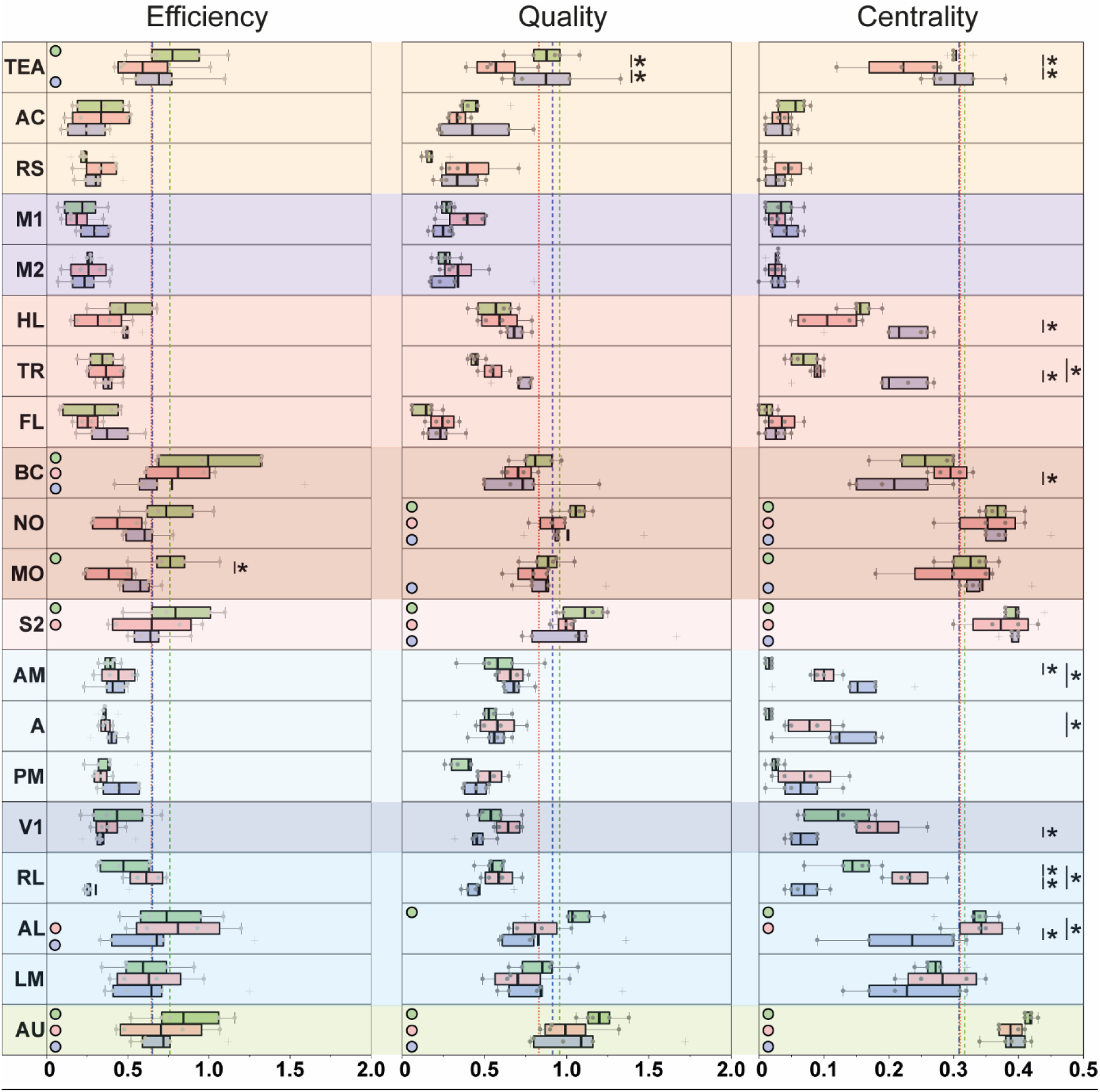
– Effects of Early Visual Deprivation and Congenital Anophthalmia on Cortical Nodal Influence Metrics in ZRDBA Mice. Cortical distributions of nodal influence metrics—namely nodal efficiency, connectivity strength, and eigenvector centrality—are presented for sighted (green), enucleated (red), and anophthalmic (blue) ZRDBA mice. Colored markers indicate cortical hubs for each metric, defined as nodes with values exceeding one standard deviation above the network mean (threshold shown as dashed lines). Boxplots display the interquartile range (25th–75th percentiles), with whiskers representing the 95% confidence interval; outliers are depicted as crosses and were retained in all statistical analyses, as no methodological or biological rationale justified their exclusion. The background color reflects the functional affiliation of each cortical area, following the functional grouping scheme described in Figure 3. Statistically significant differences between groups, based on Bonferroni-corrected tests (p ≤ 0.05), are indicated by asterisks (*).

In the visual cortex of sighted ZRDBA network, the anterolateral HVA (AL) was classified as both high-quality and core hub under sighted conditions. Although its high-quality hub status was lost in both enucleated and anophthalmic groups, its core hub designation was only lost in anophthalmic mice. Nonetheless, AL was maintained as an efficient hub in both early blindness models. Univariate analysis demonstrated that V1 centrality was significantly reduced in anophthalmic mice compared to enucleated animals (p < 0.001). Lateral and medial visual areas exhibited divergent trajectories in the anophthalmic but not in the enucleated ZRDBA mice. Centrality of RL was increased in enucleated relative to sighted controls (p = 0.010), whereas it was decreased in both AL and RL in anophthalmic compared to sighted mice (p = 0.001 and p < 0.001). In contrast, medial HVAs, A and AM, demonstrated increased nodal influence. Centrality in AM was significantly elevated in both enucleated (p = 0.016) and anophthalmic (p < 0.001) mice, while area A also showed a significant increase in centrality only in anophtalmic (p < 0.001) compared to sighted mice.

In the somatosensory and motor cortices of sighted ZRDBA network, S2 qualified as an efficient, high-quality and core hub, MO as an efficient and core hub, BC as an efficient hub and NO as a high-quality hub. In the enucleated ZRDBA mice, BC, NO, and S2 retained their classifications, whereas MO lost both efficient and core hub status. In the anophthalmic mice, BC and NO preserved all designations, while S2 and MO showed a selective loss of efficient hub classification. BC showed reduced centrality in anophthalmic compared to enucleated mice (p = 0.012). The most prominent reorganization was observed in the trunk-related area (TR), which exhibited significantly increased centrality in anophthalmic mice compared to both sighted and enucleated mice (p < 0.001). The hind limb (HL) region also showed a significant increase in centrality in anophthalmic relative to enucleated mice (p < 0.001).

In associative and auditory regions, the primary auditory cortex (AU) was the only area consistently classified as an efficient, high-quality and core hub across all visual conditions, underscoring its stable integrative role. The temporal association area (TEA) also functioned as an efficient hub in both sighted and anophthalmic ZRDBA mice. In enucleated animals, TEA exhibited significant reductions in connectivity quality (p = 0.025) and centrality (p = 0.002) compared to sighted controls. Notably, anophthalmic mice showed significantly elevated values in both metrics relative to enucleated counterparts (quality: p = 0.027; centrality: p = 0.025).

#### Differential Cortical Network Reorganization Following Neonatal Enucleation in C57BL/6J and ZRDBA Mice

Multivariate analysis (excluding anophthalmic ZRDBA mice) revealed a significant interaction between strain and visual status (Pillai’s trace: V = 0.064, F [3, 354] = 8.065, p < 0.001). Enucleation led to strain-dependent changes in global efficiency (p = 0.005) and quality metrics (p = 0.010).

Neonatal enucleation produced distinct nodal influence patterns across strains. In C57BL/6J mice, reductions were observed in primary and lateral visual cortices, while medial visual, somatosensory, and motor areas showed increased influence. In ZRDBA mice, alterations were predominantly confined to visual cortical regions. Enucleated individuals from both strains demonstrated heightened centrality in high-order visual areas, whereas anophthalmic mice exhibited diminished nodal influence in lateral HVAs and enhanced connectivity in medial HVAs. Notably, the nodal influential profile of anophthalmic ZRDBA mice more closely aligned with that of enucleated C57Bl/6J mice than with enucleated ZRDBA mice.

### Functional Network Architecture and Intermodular Connectivity Following Early Blindness Across Mouse Strains

To investigate how early blindness altered the modular architecture of the cortical connectome, we applied Louvain community detection (Blondel et al. 2008) to identify modules based on intra- and interhemispheric correlation patterns. This method captures the balance between functional segregation and integration by maximizing the modularity. To quantify the similarity between modular structures between experimental groups, we used normalized mutual information (NMI), which measures the similarity between the allocation of nodes to each community, scaled from 0 (no similarity) to 1 (identical structure). To assess the role of individual nodes within this modular framework, we computed the same three complementary metrics: participation coefficient (PC), within-module centrality (WMC), and betweenness centrality (BeC). PC quantifies a node’s connectivity across modules, identifying cross-community integrators (Guimera and Nunes Amaral 2005). WMC reflects a node’s importance within its own module, highlighting local hubs of cohesion. BeC measures how often a node lies on the shortest paths between others, indicating its role in global communication (van den Heuvel and Sporns 2011). As above, nodes with values exceeding one standard deviation above the mean plus for a given metric was classified as provincial hubs (WMC), connector nodes (PC), or bridging nodes (BeC), respectively. Together, these metrics provide a detailed view of how cortical regions contribute to intra- and intermodular connectivity in the context of congenital and early acquired blindness.

#### Neonatal Enucleation Induces Modular Reduction and Functional Reorganization of Structural Hubs in C57BL/6J Cortical Networks

In sighted C57Bl/6J mice, Louvain community detection revealed four distinct cortical intrahemispheric modules (Figure 7A). The first community (in blue) consisted solely of the associative region (RS) and had no structural hub. The second community (in yellow) included motor regions (M1, M2) and associative area (AC), anchored by M2 which served as both a provincial hub and a connector node; AC and M1 were also identified as connector nodes. The third community (in red) comprised dorsal column-related somatosensory areas (FL, HL, TR) and medial visual cortices (A, AM, PM). It was centered around TR, identified as both a provincial and bridging hub, while PM served as a connector node. The fourth community (in purple) encompassed trigeminal-related regions (BC, MO, NO), supplementary somatosensory area (S2), primary visual cortex (V1), lateral visual areas (AL, LM, RL), auditory region (AU), and associative cortex (TEA). AL and AU were identified as provincial hubs, with AU also acting as a bridging node.

**Figure 7.**
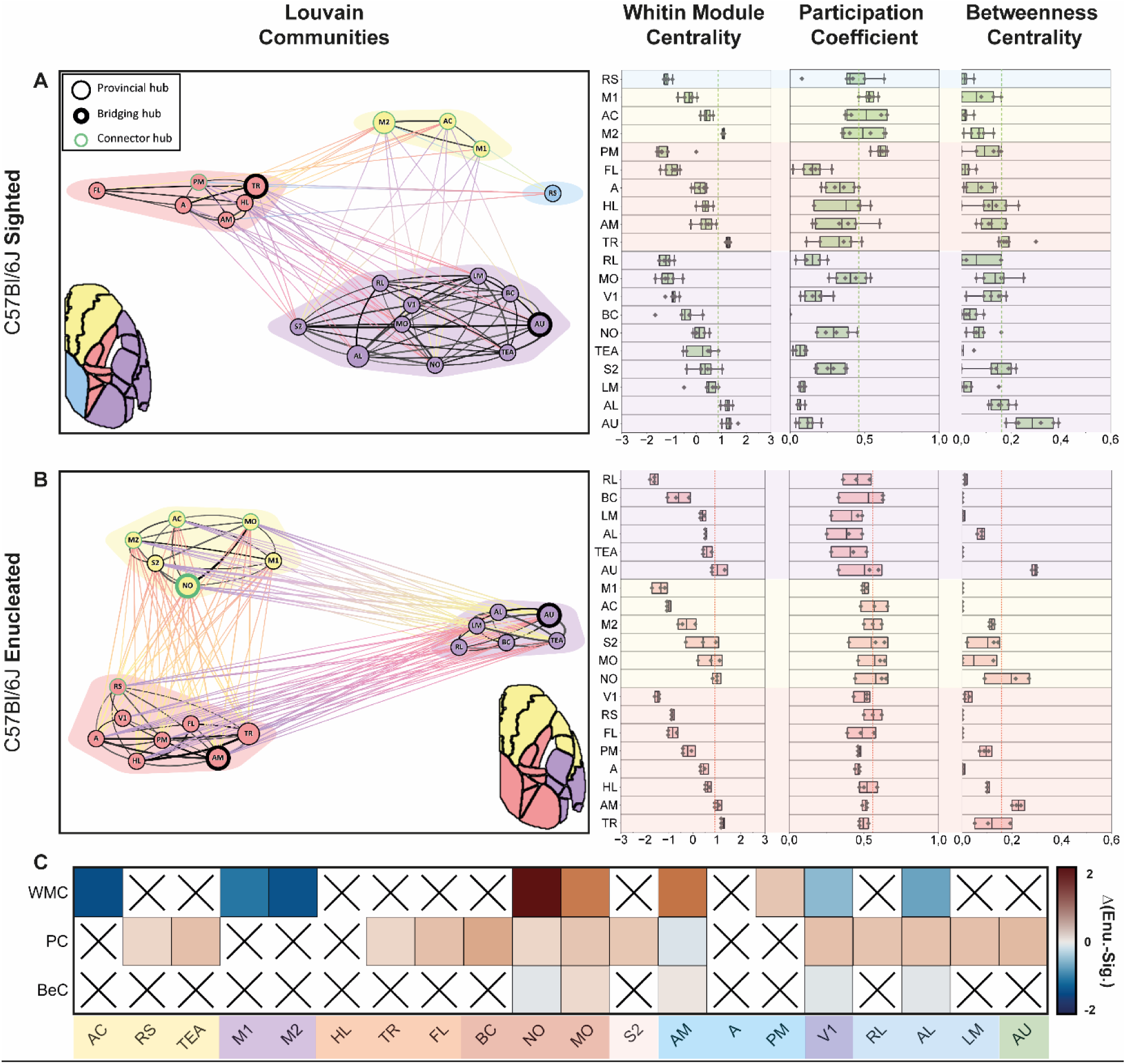
– Network Community Structure and Structural Nodal Metrics in sighted (A), enucleated (B), and group-difference mapping (C) in C57Bl/6J mice. **A.** Community structure of sighted C57Bl/6J mice obtained using Louvain modularity analysis. Community affiliations are color-coded, and the background color reflects the modular assignment of each cortical area. Provincial hubs are represented by enlarged markers, connector nodes by markers outlined in green, and bridging nodes by markers with thickened outlines. Edge thickness scales with correlation strength. Hubs are defined as nodes with values exceeding the group mean by more than one standard deviation (average + 1 STD). Structural nodal metrics—including within-module degree z-score, participation coefficient, and betweenness centrality—are shown as boxplots positioned beside the network representation. Boxplots indicate the 25th–75th percentiles, whiskers denote the 95% confidence interval, and dashed threshold lines correspond to hub-classification criteria. Outliers are displayed as crosses and were retained in the statistical analyses, as no methodological or biological justification supported their exclusion. **B.** Community structure of enucleated C57Bl/6J mice, displayed using the same modular color-coding, marker conventions, and hub definitions as in A. Background color indicates modular affiliation. Structural nodal metrics for this group are presented as corresponding boxplots, following the same statistical and graphical conventions. **C.** Heatmap depicting significant between-group differences in structural nodal metrics across cortical areas. Cells with color indicate measures exhibiting statistically significant differences between sighted and enucleated mice, based on MANOVA followed by Bonferroni-corrected post-hoc comparisons (p ≤ 0.05). Crosses denote metrics for which no significant difference was detected.

In enucleated C57Bl/6J mice (Figure 7B), the cortical network reorganized into three communities. The first community (in yellow) included somatosensory regions (MO, NO, S2), associative area (AC), and motor regions (M1, M2). NO emerged as a provincial hub, a connector, and a bridging node, highlighting its prominent role in local and intermodular coordination. AC, MO, and M2 were also identified as connector nodes. The second community (in red) included dorsal column-related somatosensory areas (FL, HL, TR), medial visual cortices (A, AM, PM), primary visual cortex (V1), and associative region (RS). AM and TR were identified as provincial hubs, with AM also serving as a bridging node and RS as a connector node. The third community (in purple) was composed of the somatosensory region (BC), lateral visual cortices (AL, LM, RL), auditory area (AU), and the associative cortex (TEA). AU was identified as both a provincial hub and a bridging node.

##### Cortical Community Reorganization and Node Role Shifts Following Enucleation in C57Bl/6J Mice

Overall, NMI between cortical networks communities of sighted and enucleated mice reached 57.4%. Multivariate analysis revealed significant effects of enucleation on nodal structural metrics, including within-module centrality, participation coefficients, and betweenness centrality, with a strong interaction between visual status and cortical region (Pillai’s trace: V = 1.274, F [57, 420] = 5.436, p < 0.001). Nearly all regions were significantly affected (except for HL), reflecting a redistribution of local and intermodular influence and a profound reorganization of cortical network.

Univariate statistical analysis further highlighted significant nodal changes within the structural hubs of the enucleated C57Bl/6J mice cortical network. Among provincial hubs, AU and TR remained stable, whereas M2 (p < 0.001) and AL (p = 0.003) lost their hub designation. Conversely, AM (p = 0.007) and NO (p < 0.001) emerged as new provincial hubs, indicating an increased involvement of somatosensory and medial visual areas in cortical processing. Connector node dynamics followed a similar trend, with AC and M2 maintaining their roles, while MO (p = 0.022), NO (p < 0.001), and RS (p = 0.026) were newly classified as connector nodes. Although M1 and PM lost their connector status, their changes in participation coefficients did not reach statistical significance. Modifications in bridging nodes further illustrated the network’s reorganization. AU preserved its bridging role, whereas TR showed a nonsignificant increase in betweenness centrality but lost its bridging node status. In contrast, AM (p = 0.004) and NO (p < 0.001) no longer served as bridging nodes due to significant reductions in betweenness centrality. These results point to a shift in long-range cortical communication, favoring distributed integration over centralized bottlenecks. To summarize, nodal structural reorganization supported by statistical analysis included a significant loss of provincial hub status for AL and M2, while AM and NO significantly emerged as new provincial hubs. Additionally, MO, NO, and RS gained connector node classification, each driven by statistically significant increases in participation coefficients. AM and NO also transitioned into bridging node roles, supported by significant elevations in betweenness centrality.

###### Visual Deprivation Models Reshape Cortical Architecture Through Modular Reduction and Nodal Structural Reclassification in ZRDBA Mice

In sighted ZRDBA mice (Figure 8A), Louvain analysis identified four distinct cortical communities. An anterior community (in yellow) was composed of AC and M2 and both were labelled as connector nodes. A second community (in blue) integrated dorsal column related somatosensory (FL, HL, TR) and motor (M1) areas. In this community, TR was identified as both connector and bridging node, while HL was also considered as a bridging node. A third community (in purple) incorporated trigeminal-related (BC, MO, NO) and supplemental somatosensory (S2), lateral (AL, LM, RL) and primary visual (V1) and auditory (AU) areas. In terms of nodal structure, AU and S2 functioned as provincial hubs, while AU was also identified as a bridging node. Additionally, AL was also considered as a bridging node. Finally, a fourth community (in red) was composed of associative (RS) and medial visual (A, AM, PM) areas. In this community, AM was a provincial node, while PM was a bridging node.

**Figure 8.**
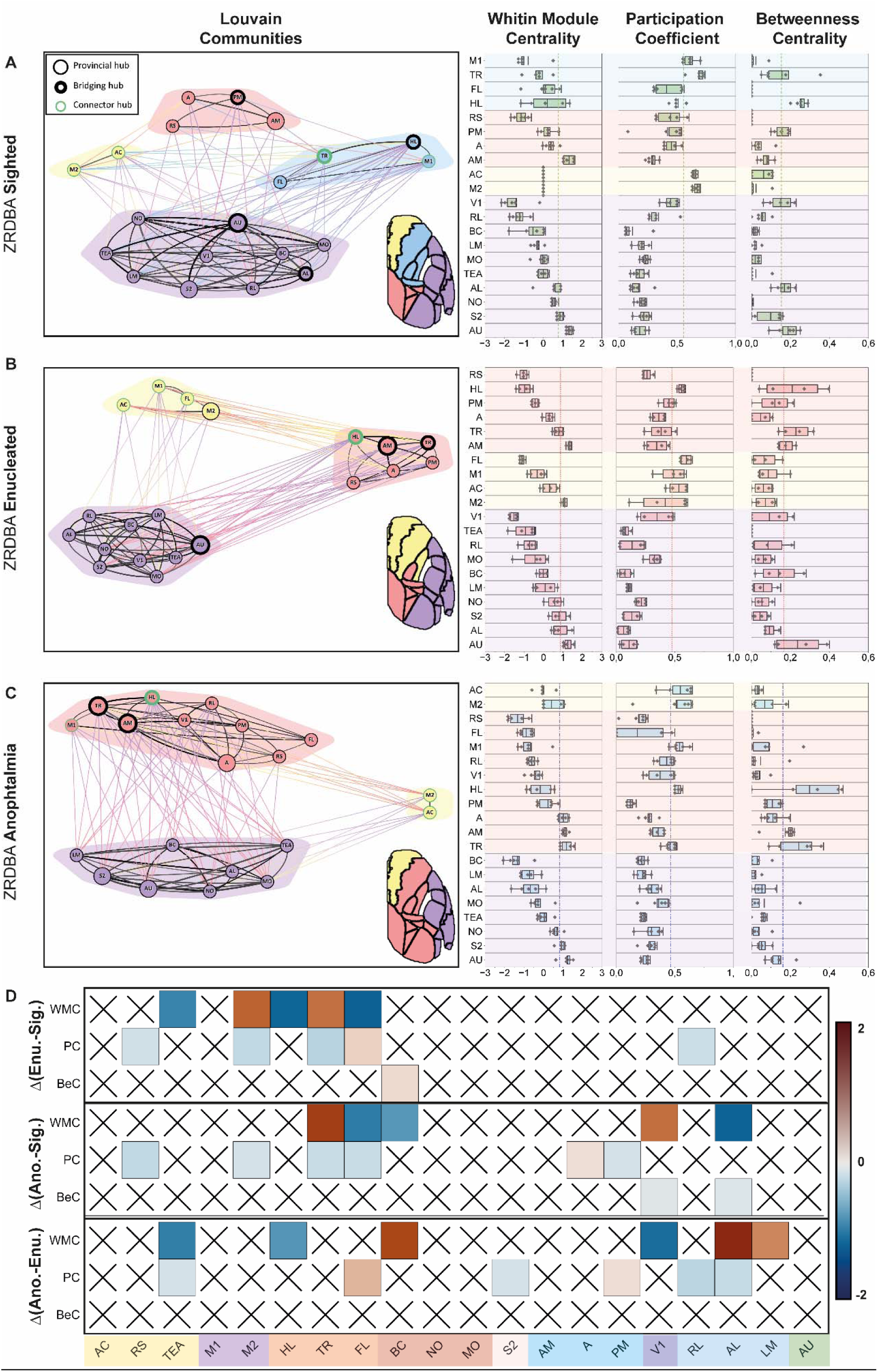
– Network Community Structure and Structural Nodal Metrics in sighted (A), enucleated (B), anophthalmic (C), and group-difference mapping (D) in ZRDBA mice. **A.** Community structure of sighted ZRDBA mice obtained using Louvain modularity analysis. Community affiliations are color-coded, and the background color reflects the modular assignment for each cortical area. Provincial hubs appear as enlarged markers, connector nodes as markers outlined in green, and bridging nodes as markers with thickened outlines. Edge thickness is proportional to connection strength. Hubs are defined as nodes whose values exceed the group mean by more than one standard deviation (average + 1 STD). Structural nodal metrics—including within-module degree z-score, participation coefficient, and betweenness centrality—are displayed as boxplots positioned beside the network representation. Boxplots show the interquartile range (25th–75th percentiles), whiskers represent the 95% confidence interval, and threshold dashed lines indicate the criteria for structural hub classification. Outliers are represented by crosses and were retained in the analyses, as no methodological or biological rationale justified their exclusion. **B.** Community structure of enucleated ZRDBA mice, using the same modular color scheme and hub definitions as in **A.** Background color indicates modular affiliation. Structural nodal metrics for this group are shown as corresponding boxplots, with identical conventions for thresholds, whiskers, and outliers. **C.** Community structure of anophthalmic ZRDBA mice, following the same analytical and graphical conventions as in **A.** and **B**. Background color reflects modular assignment. Boxplots for within-module degree z-score, participation coefficient, and betweenness centrality are displayed beside the network, with the same threshold lines and outlier representation. **D.** Heatmap summarizing significant between-group differences in structural nodal metrics across cortical areas for the three ZRDBA conditions (sighted, enucleated, anophthalmic). Colored cells indicate metrics showing statistically significant differences based on MANOVA followed by Bonferroni-corrected post-hoc comparisons (p ≤ 0.05). Crosses denote cortical regions and metrics for which no significant difference was detected.

In the enucleated ZRDBA mice (Figure 8B), the cortical network was restructured into three communities. A first community (in yellow), which incorporated the somatomotor (FL, M1, M2) areas, in which M2 qualified as a provincial hub. Additionally, AC, FL and M1 were considered as connector nodes. A second community (in red) contained associative (RS), dorsal column-related somatosensory (HL, TR) and medial visual (A, AM, PM) areas. AM had both a provincial hub and a bridging node function. HL and TR were also considered as bridging nodes, additionally, HL was also qualified as a connector node. A third community (in purple) encompassed associative (TEA), auditory (AU), trigeminal-related (BC, MO, NO) and supplemental somatosensory (S2) and lateral (AL, LM, RL) and primary visual (V1). AU acted as both a provincial hub and a bridging node.

In anophthalmic ZRDBA mice (Figure 8C), we observed three distinct cortical communities. A first community (in yellow), having a structure similar as to sighted mice was composed of AC and M2, which were both connector nodes. A second community (in purple) comprised associative (TEA), auditory (AU), trigeminal (BC, MO, NO) and supplemental somatosensory (S2) and lateral visual (AL and LM) areas with AU and S2 as provincial hubs. A third community (in red) was composed by associative (RS), motor (M1), dorsal column related somatosensory (FL, HL and TR) and visual (A, AM, PM, RL and V1) areas. AM served as a provincial hub alongside HL and TR. Together with TR, both AM and HL also acted as bridging nodes. Additionally, M1 and HL functioned as connector hubs.

##### Cortical Community Structure and Nodal Function Modulation Following Early Blindness Across Distinct Visual Conditions in ZRDBA mice

Comparative analysis of cortical network communities revealed moderate overlap in structural organization across visual conditions. In the ZRDBA mice, NMI values were 79.2% (sighted vs. enucleated), 64.1% (sighted vs. anophthalmic), and 56.2% (anophthalmic vs. enucleated). Multivariate analysis demonstrated significant effects of enucleation on nodal metrics including within-module centrality, participation coefficients, and betweenness centrality, with a robust interaction between visual status and cortical region (Pillai’s trace: V = 1.079, F [114, 660] = 3.254, p < 0.001). Early blindness significantly impacted associative (RS, TEA), motor (M2), somatosensory (BC, FL, HL, TR), visual (A, AL, PM, RL, V1), and auditory (AU) regions.

Univariate analysis comparing sighted and enucleated ZRDBA mice revealed notable shifts in nodal structural function across cortical communities. Among the identified provincial hubs, AU and AM maintained stable roles across conditions. M2 emerged as a new provincial hub, marked by a significant increase in WMC (p = 0.001). Conversely, S2 lost its hub status despite exhibiting no significant change in WMC. For the connector nodes, AC and M1 retained their roles, with participation coefficients remaining unchanged. In contrast, M2 and TR lost their connector node classification, exhibiting a significant decrease in the participation coefficient (both p < 0.001). Additionally, FL and HL gained connector node status, with FL showing a significant increase in participation coefficient (p = 0.010), whereas HL’s rise did not reach statistical significance. For the bridging nodes, AU, HL and TR maintained their bridging roles, with betweenness centrality showing no significant changes. PM and AL, however, lost their classification as bridging nodes, though their betweenness centrality decreases were not statistically significant. AM, in contrast, gained bridging node status, yet this shift was not accompanied by a significant betweenness centrality increase. To summarize, nodal structural reorganization supported by statistical analysis between sighted and enucleated mice included M2’s dual shift—emerging as a provincial hub while losing its connector node status. TR also lost connector classification due to a significant decline in participation coefficients, whereas FL gained connector node status, driven by a significant increase in participation coefficients.

Univariate analysis comparing sighted and anophthalmic ZRDBA mice revealed key shifts in nodal roles within cortical communities. Among provincial hubs, AM, AU and S2 maintained their hub status, showing no significant changes in WMC, while TR and A gained provincial hub classification, though only the WMC of TR was significantly increased (p < 0.001). Changes in connector nodes followed a similar trend, with AC, M1, and M2 retaining their roles, as their participation coefficients remained unchanged (except for M2; p = 0.025). In contrast, TR lost its connector node classification, through a participation coefficient significant decrease (p < 0.001). In addition, HL gained the connector node status, yet its participation coefficient increase did not reach statistical significance. HL and TR consistently retained their roles as bridging nodes, with no significant changes of their betweenness centrality score. AU and PM lost their bridging status, although declines in betweenness centrality were not statistically significant. AL also lost its classification, accompanied by a significant reduction in betweenness centrality (p = 0.033). AM gained bridging node status, though this reclassification was not supported by a significant increase in betweenness centrality.

To summarize, statistically supported nodal structural reorganization between sighted and anophtalmic mice included TR’s emergence as a provincial hub alongside its loss of connector status, and AL’s loss of bridging node classification driven by a significant decrease in betweenness centrality.

Univariate analysis comparing anophthalmic and enucleated ZRDBA mice revealed key differences in nodal roles across cortical communities. Among provincial hubs, AU and AM maintained their classification, with no significant changes in WMC. Conversely, A, TR and S2 lost their provincial hub designation, though their WMC reductions were not statistically significant, while M2 gained provincial hub status without significant decrease of WMC. Having stable participation coefficients, AC, M1, and HL remained connector nodes. In contrast, M2 lost its connector node status without a significant decrease in participation coefficients, while FL emerged as a new connector node with a statistically significant increase in participation coefficient (p < 0.001). For the bridging nodes, AM, HL and TR maintained their function with no significant alterations in betweenness centrality, whereas AU gained the classification, though its increase in betweenness centrality was not significant. Overall, the only statistically supported structural hub alteration between anophthalmia and enucleation was the emergence of FL as a connector node, driven by increased participation.

###### Visual Deprivation and Genetic Background Synergistically Reshape Cortical Network Modularity

Enucleated C57BL/6J and both early blind ZRDBA mice showed reduced cortical modularity, with the number of communities dropping from four to three, combined with increased intermodular connectivity. Between sighted C57BL/6J and ZRDBA mice NMI reached 77.6% but declined to 49.8% between their enucleated counterparts, indicating greater structural divergence after early enucleation. Multivariate analysis revealed a significant interaction between strain and visual status on network topology (Pillai’s trace: V = 0.156, F [3, 354] = 21.780, p < 0.001). Although WMC and betweenness centrality did not differ significantly across strains, participation coefficient values were significantly elevated in C57BL/6J mice (p < 0.001), suggesting stronger intermodular integration compared to ZRDBA.

## Discussion

### Alteration of the infraslow oscillation activity

Our findings demonstrate that early visual deprivation altered the spectral power of cortical activity, particularly in the infraslow oscillation (ISO) range, across both mouse strains studied. In C57Bl/6J mice, neonatal enucleation led to a generalized increase in ISO power, most prominently in associative, dorsal column-related somatosensory and medial visual cortices. In contrast, enucleated ZRDBA mice showed more limited enhancements, primarily in auditory and trigeminal-related somatosensory cortices. These strain-specific patterns suggest distinct forms of large-scale cortical reorganization in response to early loss of vision.

Prior studies have reported that sensory deprivation alters ISOs in both mice and humans (Burton *et al*. 2014; Wang *et al*. 2014; Heine *et al*. 2015; Kraft et al. 2017). ISOs are recognized as markers of intrinsic large-scale cortical connectivity and excitability (Vanhatalo et al. 2004; Fox et al. 2005; Fox et al. 2006; Power et al. 2011; Mitra et al. 2014; Sforazzini et al. 2014; Mitra et al. 2015). Our data reveal that the spatial pattern of ISO power increases in enucleated C57BL/6J mice closely resembles that observed in anophthalmic ZRDBA mice yet differs from the profile found in enucleated ZRDBA animals. This divergence likely reflects a genetic contribution to cortical plasticity. Such an interpretation is consistent with previously documented strain-dependent differences in cortical plasticity between C57BL/6 and DBA mice (Heimel *et al*. 2008), as well as with our earlier findings showing strain-specific variations in the size and developmental trajectory of the visual cortex following either enucleation or anophthalmia in C57BL/6J and ZRDCT/An mice (Masse et al. 2014). Notably, ZRDBA mice originate from a cross between DBA and ZRDCT/An strains, further supporting this genetic influence. Moreover, within the ZRDBA strain itself, we observed distinct ISO power profiles when comparing anophthalmic and enucleated animals. This intra-strain difference underscores the respective roles of prenatal and postnatal afferent activity in shaping cortical functional development (Masse *et al*. 2014).

Human studies bear both similarities and differences with what we observe here in blind mice. In congenitally blind individuals, reorganization affects visual, auditory, frontal, and retrosplenial regions (Lubinus et al. 2021), consistent with the territories modulated in our murine models. However, humans show more extensive spectral changes, including increased theta and beta power and reduced alpha activity (Noebels et al. 1978; Kriegseis et al. 2006; Schepers et al. 2012; Hawellek et al. 2013; Schubert et al. 2015; Gudi-Mindermann et al. 2018; Rimmele et al. 2019). In contrast, our recordings revealed changes predominantly below 4 Hz, concentrated in the infraslow and delta bands. Although some signal extended into theta and alpha ranges, no consistent group differences emerged beyond 4 Hz. This discrepancy may be attributed to species-specific plasticity profiles, but methodological limitations such as the slow kinetics of calcium indicators (GCaMP6s) likely contributed as well. Additionally, our data does not capture the full spectral range, notably excluding beta and gamma oscillations.

### Alteration of interhemispheric functional connectivity

Our analysis revealed strain-specific patterns of cortical network reorganizations following early visual deprivation. In C57Bl/6J enucleated mice, we observed widespread increases in both heterotopic and homotopic interhemispheric connectivity of associative, auditory, and visual regions, suggesting wide-ranging consequences of visual deprivation, well beyond the realm of visual cortical areas. ZRDBA enucleated mice exhibited subtler changes, with stable homotopic connectivity and selective heterotopic enhancements, particularly involving medial higher visual areas. In contrast, ZRDBA anophthalmic mice showed reduced homotopic connectivity in associative and motor cortices, while maintaining similar heterotopic enhancements of medial HVAs as observed in enucleated ZRDBA mice.

The widespread increases in homotopic and heterotopic interhemispheric connectivity in C57Bl/6J enucleated mice, notably involving associative and auditory mirror findings in congenitally blind humans, in whom non-visual networks exhibit enhanced cross-hemispheric integration (Bedny *et al*. 2011; Burton *et al*. 2014). However, increased interhemispheric functional connectivity between visual cortices diverges from human data (Liu *et al*. 2007; Yu *et al*. 2008; Watkins *et al*. 2012; Butt et al. 2013; Burton *et al*. 2014; Bauer *et al*. 2017), potentially reflecting species-specific effects.

structural MRI studies report thinning of the posterior corpus callosum in early blind individuals, which supports reduced visual interhemispheric transfer (Ptito et al. 2008; Lepore et al. 2010), and increases in the anterior and mid-body regions suggesting an enhanced integration across frontal, somatosensory, and auditory areas (Ptito *et al*. 2008; Tomaiuolo et al. 2014). However, other reports found no callosal changes in early blind or anophthalmic individuals (Bock et al. 2013), indicating variability across cohorts. In contrast, animal models consistently show disrupted development of visual callosal projections following deprivation paradigms such as dark rearing, eyelid sutures, or optic chiasm sectioning (Lund and Mitchell 1979; Innocenti et al. 1985; Frost and Moy 1989; Boire et al. 1995). In rodents, enucleation and anophthalmia often produce broader callosal neuron distributions, extending beyond the typical boundaries of primary and extrastriate cortices (Olavarria and van Sluyters 1984; Rhoades *et al*. 1984; Olavarria et al. 1987; Olavarria *et al*. 1988; Olavarria and Li 1995).

The targeted reorganization seen in ZRDBA enucleated mice, marked by stable homotopic connectivity and selective heterotopic increases, resembles the patchwork pattern of interhemispheric connectivity modifications described in other human studies (Qin et al. 2013; Bridge and Watkins 2019), which highlight localized adaptations in the absence of vision.

### Cortical Connectivity Modifications Following Early Visual Deprivation

Resting-state functional connectivity analyses revealed similar intrahemispheric cortical network in the intact and enucleated ZRDBA mice. However more substantial differences were observed in the anophthalmic mice. This suggests a greater impact of prenatal than postnatal activity of the development of cortical functional. This contrasts with the significant differences we observed between the intact and enucleated C57BL6 mice which suggest that, in this strain, prenatal retinal activity does interfere with cortical development. This is could be related to the superior visual cortex plasticity of the C57BL6 compared to the DBA mice (Heimel *et al*. 2008).

Enucleated ZRDBA mice retain residual prenatal retinal activity but fail to show substantial reorganization, suggesting this limited activity either lacks the strength to override pre-established spontaneous wiring or sufficiently preserves thalamocortical integrity. In contrast, complete absence of afferent retinal input in anophthalmic ZRDBA animals has a more pronounced effect on cortical development.

These results underscore a multifactorial influence, genetic background, developmental timing, and prenatal spontaneous activity, in shaping cortical reorganization trajectories. The C57Bl/6J strain, owing to its high plastic potential, enucleation reshapes cortical networks in a manner that converges toward genetically anchored configurations like those observed in congenitally blind ZRDBA mice.

These strain-specific and developmental differences have translational relevance, echoing findings on genetically mediated plasticity (Kaschube et al. 2002; Missitzi et al. 2011) and contributing to the understanding of variability in human responses to sensory deprivation. In anophthalmic mice, reallocation of subcortical projections might predominate (Rhoades *et al*. 1985; Chabot et al. 2008), whereas enucleation might favor cortical reorganization involving associative and higher-order visual areas.

### Area-Specific Plasticity: Insights From V1

Changes in connectivity were not uniform across the cortex. Notably, V1 in enucleated C57Bl/6J mice exhibited diminished intrahemispheric connectivity with somatosensory and auditory cortices, but increased connectivity with associative regions, such as the retrosplenial cortex. This reorganization echoes findings in early blind humans, where V1 becomes more tightly coupled to multimodal association areas and less connected to non-visual primary sensory and motor cortices (Burton *et al*. 2014; Bauer *et al*. 2017; Abboud and Cohen 2019).

This shift may be driven by V1’s newly acquired responsiveness to non-visual stimuli. Functional activation of the visual cortex by tactile and auditory inputs has been consistently reported in blind subjects (Hyvarinen et al. 1981; Roder et al. 1996; Sadato *et al*. 1996; Cohen et al. 1997; Kujala et al. 1997; Sadato et al. 1998; Weeks et al. 2000) and is often interpreted as evidence of enhanced crossmodal connectivity (Wittenberg et al. 2004; Petrus *et al*. 2014; Petrus *et al*. 2015).

Other studies, however, document a decrease in connectivity between V1 and spared sensory cortices (Liu *et al*. 2007; Yu *et al*. 2008; Shu et al. 2009), revealing the complexity of crossmodal plasticity and its dependence on developmental timing and network architecture.

### Higher Visual Areas as Dynamic Network Hubs

In our study, one of the most compelling observations was the emergence of medial HVAs, particularly area AM, as influential and structural hub within the reorganized cortical network of enucleated C57Bl/6J mice. These regions exhibited increased functional connectivity with non-visual sensory cortices, suggesting that medial HVAs play a central integrative role following early sensory loss. In contrast, primary and lateral visual cortices showed reduced nodal influence, highlighting a functional divergence among HVAs that may reflect differential specialization in crossmodal processing.

Anatomical studies in sighted mice support our findings, demonstrating that HVAs, particularly PM, LM, and AM, as well as the associative retrosplenial area, provide robust top-down projections to V1 (Oh *et al*. 2014; Zingg *et al*. 2014; Morimoto et al. 2021). Additionally, anatomical studies in visually deprived mice confirm that these projections remain structurally intact and may be functionally strengthened following sensory loss. This would be in agreement with the connectivity constrained experience dependency hypothesis (Saccone *et al*. 2024). In fact, it was demonstrated that HVAs maintain top-down projections to V1 in enucleated mice (Laramee et al. 2011; Gilissen and Arckens 2021) and that V1 receives direct inputs from auditory and somatosensory in both enucleated and anophtalmic mice (Charbonneau *et al*. 2012), suggesting that crossmodal integration may rely on pre-existing corticocortical pathways.

Human studies support this organizational framework across both sighted and blind individuals. In sighted subjects, structural imaging has revealed intrinsic anatomical connectivity between visual and non-visual sensory cortices, supporting the potential for multisensory integration even in the presence of intact vision (Foxe and Schroeder 2005; Macaluso and Driver 2005; Macaluso 2006; Ricciardi, Bonino, et al. 2014). In blind individuals, this system undergoes experience-dependent reorganization, with increased structural and functional coupling between occipital areas and auditory or tactile regions (Noppeney et al. 2005; Collignon *et al*. 2007; Collignon *et al*. 2009; Klinge et al. 2010; Bedny *et al*. 2011; Collignon et al. 2011; Collignon et al. 2013). Activation of the occipital cortex during tactile and auditory tasks appears to be driven primarily by top-down feedback from dorsal and ventral HVAs (Ricciardi, Tozzi, et al. 2014; Murphy *et al*. 2016), while direct connections from primary sensory cortices to V1 tend to diminish (Liu *et al*. 2007; Ortiz-Terán et al. 2017). These pathways suggest that HVAs and associative regions are strategically positioned to influence V1 activity, even in the absence of visual input, reinforcing their potential importance in cross-modal recruitment of V1.

### Retrosplenial Cortex: A Consistent Integrative Node

Across all models, the retrosplenial cortex (RS) consistently gained influence within the functional network. Long considered a hub for spatial processing, the RS is increasingly recognized for its role in multisensory and unimodal integration (Hindley et al. 2014; Fournier et al. 2020; Alexander et al. 2023). Its elevated centrality in enucleated mice suggests that associative regions play a crucial role in network dynamics after sensory loss. This supports the hypothesis that existing cortico-cortical projections are recruited during crossmodal plasticity (Smolders et al. 2016; Barnes et al. 2022), with RS serving as a bridge between deprived and active sensory regions. These results also support that the connectivity constrained hypothesis (Saccone *et al*. 2024) also applies to our murine models of visual deprivation.

### Functional Reorganization Through Reweighted Pre-Existing Circuits

The changes in functional connectivity we show here are likely not the reflections of the quite modest alterations in anatomical connectivity as shown by tract tracing studies in enucleated and anophthalmic mice (Laramee *et al*. 2011; Charbonneau *et al*. 2012; Laramée *et al*. 2014; Laramée et al. 2016). There is accumulating evidence that selective potentiation of pre-existing circuits through Hebbian and homeostatic mechanisms (Lee and Whitt 2015; Ewall et al. 2021; Makin and Krakauer 2023), notably descending multisensory projections and latent intracortical pathways (Meng et al. 2015; Petrus *et al*. 2015).

Following visual deprivation the preserved sensory modalities exhibit enhanced local inhibition (Petrus et al. 2014), which sharpens unimodal processing and thwarts cross-modal interference. Simultaneously, the deprived visual cortex undergoes potentiation of corticocortical inputs (Petrus *et al*. 2015; Lee 2022), increased excitatory drive. Experimental evidence has shown that blindness induces synaptic scaling, alterations in excitation–inhibition balance, and heightened cortical excitability (Keck et al. 2013; Gainey and Feldman 2017). In rodents, these adaptations include potentiated thalamocortical and intracortical connectivity in spared sensory pathways (Gao et al. 2010; Gao et al. 2014; Petrus *et al*. 2014; Petrus *et al*. 2015; Ewall *et al*. 2021), in agreement with reallocation of cortical processing. Moreover, our results further documents that such changes shape both local circuitry and whole-brain functional connectivity (Lambo and Turrigiano 2013; Hellyer et al. 2016; Rocha et al. 2018).

## Conclusion

In conclusion, our study demonstrates that neonatal sensory deprivation profoundly alters cortical connectivity, with distinct patterns of reorganization depending on the timing of sensory loss and genetic background. The differential plasticity observed between C57Bl/6J and ZRDBA mice highlight the interplay between genetic factors and sensory experience in shaping cortical networks. By elucidating the mechanisms underlying crossmodal reorganization and the role of higher visual areas, our findings provide novel insights into the adaptive capacity of the brain following sensory deprivation.

While our study provides important insights into the large-scale cortical reorganization following neonatal sensory deprivation, a more granular understanding of the underlying cellular mechanisms is needed. Specifically, the role of distinct neuronal populations, excitatory and inhibitory, in these processes remains largely unexplored. Investigating how sensory deprivation alters the activity and connectivity of specific excitatory populations, such as pyramidal neurons, and inhibitory populations, such as interneurons, will be pivotal in elucidating the dynamics of cortical reorganization. Furthermore, it is crucial to explore how these alterations affect synaptic plasticity, neurotransmitter release, and the fine-tuned balance between excitation and inhibition that is critical for maintaining network stability.

## REFERENCES

Abboud S, Cohen L. 2019. Distinctive Interaction Between Cognitive Networks and the Visual Cortex in Early Blind Individuals. Cereb Cortex. 29:4725–4742.

Ackman JB, Burbridge TJ, Crair MC. 2012. Retinal waves coordinate patterned activity throughout the developing visual system. Nature. 490:219–225.

Alexander AS, Place R, Starrett MJ, Chrastil ER, Nitz DA. 2023. Rethinking retrosplenial cortex: Perspectives and predictions. Neuron. 111:150–175.

Amedi A, Floel A, Knecht S, Zohary E, Cohen LG. 2004. Transcranial magnetic stimulation of the occipital pole interferes with verbal processing in blind subjects. Nat Neurosci. 7:1266–1270.

Andelin AK, Olavarria JF, Fine I, Taber EN, Schwartz D, Kroenke CD, Stevens AA. 2018. The Effect of Onset Age of Visual Deprivation on Visual Cortex Surface Area Across-Species. Cerebral Cortex. 29:4321–4333.

Andermann ML, Kerlin AM, Roumis DK, Glickfeld LL, Reid RC. 2011. Functional Specialization of Mouse Higher Visual Cortical Areas. Neuron. 72:10.1016/j.neuron.2011.1011.1013.

Anton-Bolanos N, Sempere-Ferrandez A, Guillamon-Vivancos T, Martini FJ, Perez-Saiz L, Gezelius H, Filipchuk A, Valdeolmillos M, Lopez-Bendito G. 2019. Prenatal activity from thalamic neurons governs the emergence of functional cortical maps in mice. Science. 364:987–990.

Barnes SJ, Keller GB, Keck T. 2022. Homeostatic regulation through strengthening of neuronal network-correlated synaptic inputs. eLife. 11:e81958.

Bauer CM, Hirsch GV, Zajac L, Koo BB, Collignon O, Merabet LB. 2017. Multimodal MR-imaging reveals large-scale structural and functional connectivity changes in profound early blindness. PLoS One. 12:e0173064.

Bedny M, Pascual-Leone A, Dodell-Feder D, Fedorenko E, Saxe R. 2011. Language processing in the occipital cortex of congenitally blind adults. Proc Natl Acad Sci U S A. 108:4429–4434.

Blankenship AG, Feller MB. 2010. Mechanisms underlying spontaneous patterned activity in developing neural circuits. Nat Rev Neurosci. 11:18–29.

Blondel VD, Guillaume J-L, Lambiotte R, Lefebvre E. 2008. Fast unfolding of communities in large networks. Journal of Statistical Mechanics: Theory and Experiment. 2008:P10008.

Bock AS, Saenz M, Tungaraza R, Boynton GM, Bridge H, Fine I. 2013. Visual callosal topography in the absence of retinal input. Neuroimage. 81:325–334.

Boire D, Morris R, Ptito M, Lepore F, Frost DO. 1995. Effects of neonatal splitting of the optic chiasm on the development of feline visual callosal connections. Exp Brain Res. 104:275–286.

Bonaventure N, Karli P. 1968. [Appearance in the visual cortex of evoked potentials of auditive origin in mice deprived of photoreceptors]. J Physiol (Paris). 60 Suppl 2:407.

Bridge H, Watkins KE. 2019. Structural and functional brain reorganisation due to blindness: The special case of bilateral congenital anophthalmia. Neurosci Biobehav Rev. 107:765–774.

Burton H, Snyder AZ, Raichle ME. 2014. Resting state functional connectivity in early blind humans. Front Syst Neurosci. 8:51.

Butt OH, Benson NC, Datta R, Aguirre GK. 2013. The fine-scale functional correlation of striate cortex in sighted and blind people. J Neurosci. 33:16209–16219.

Cang J, Rentería RC, Kaneko M, Liu X, Copenhagen DR, Stryker MP. 2005. Development of precise maps in visual cortex requires patterned spontaneous activity in the retina. Neuron. 48:797–809.

Chabot N, Charbonneau V, Laramee ME, Tremblay R, Boire D, Bronchti G. 2008. Subcortical auditory input to the primary visual cortex in anophthalmic mice. Neurosci Lett. 433:129–134.

Chabot N, Robert S, Tremblay R, Miceli D, Boire D, Bronchti G. 2007. Audition differently activates the visual system in neonatally enucleated mice compared with anophthalmic mutants. European Journal of Neuroscience. 26:2334–2348.

Chan KY, Jang MJ, Yoo BB, Greenbaum A, Ravi N, Wu WL, Sanchez-Guardado L, Lois C, Mazmanian SK, Deverman BE, Gradinaru V. 2017. Engineered AAVs for efficient noninvasive gene delivery to the central and peripheral nervous systems. Nat Neurosci. 20:1172–1179.

Charbonneau V, Laramee ME, Boucher V, Bronchti G, Boire D. 2012. Cortical and subcortical projections to primary visual cortex in anophthalmic, enucleated and sighted mice. Eur J Neurosci. 36:2949–2963.

Chase HB. 1942. Studies on an Anophthalmic Strain of Mice. III. Results of Crosses with Other Strains. Genetics. 27:339–348.

Chase HB. 1944. Studies on an Anophthalmic Strain of Mice. IV. a Second Major Gene for Anophthalmia. Genetics. 29:264–269.

Chase HB, Chase EB. 1941. Studies on an anophthalmic strain of mice I. Embryology of the eye region. Journal of Morphology. 68:279–301.

Cohen LG, Celnik P, Pascual-Leone A, Corwell B, Falz L, Dambrosia J, Honda M, Sadato N, Gerloff C, Catala MD, Hallett M. 1997. Functional relevance of cross-modal plasticity in blind humans. Nature. 389:180–183.

Collignon O, Davare M, Olivier E, De Volder AG. 2009. Reorganisation of the right occipito-parietal stream for auditory spatial processing in early blind humans. A transcranial magnetic stimulation study. Brain Topogr. 21:232–240.

Collignon O, Dormal G, Albouy G, Vandewalle G, Voss P, Phillips C, Lepore F. 2013. Impact of blindness onset on the functional organization and the connectivity of the occipital cortex. Brain. 136:2769–2783.

Collignon O, Lassonde M, Lepore F, Bastien D, Veraart C. 2007. Functional cerebral reorganization for auditory spatial processing and auditory substitution of vision in early blind subjects. Cereb Cortex. 17:457–465.

Collignon O, Vandewalle G, Voss P, Albouy G, Charbonneau G, Lassonde M, Lepore F. 2011. Functional specialization for auditory-spatial processing in the occipital cortex of congenitally blind humans. Proc Natl Acad Sci U S A. 108:4435–4440.

Colonnese MT. 2014. Rapid developmental emergence of stable depolarization during wakefulness by inhibitory balancing of cortical network excitability. J Neurosci. 34:5477–5485.

D’Souza RD, Wang Q, Ji W, Meier AM, Kennedy H, Knoblauch K, Burkhalter A. 2022. Hierarchical and nonhierarchical features of the mouse visual cortical network. Nat Commun. 13:503.

Dupont E, Hanganu IL, Kilb W, Hirsch S, Luhmann HJ. 2006. Rapid developmental switch in the mechanisms driving early cortical columnar networks. Nature. 439:79–83.

Ewall G, Parkins S, Lin A, Jaoui Y, Lee H-K. 2021. Cortical and Subcortical Circuits for Cross-Modal Plasticity Induced by Loss of Vision. Frontiers in Neural Circuits. 15.

Finneran DJ, Njoku IP, Flores-Pazarin D, Ranabothu MR, Nash KR, Morgan D, Gordon MN. 2021. Toward Development of Neuron Specific Transduction After Systemic Delivery of Viral Vectors. Frontiers in Neurology. 12.

Fournier DI, Monasch RR, Bucci DJ, Todd TP. 2020. Retrosplenial cortex damage impairs unimodal sensory preconditioning. Behav Neurosci. 134:198–207.

Fox MD, Corbetta M, Snyder AZ, Vincent JL, Raichle ME. 2006. Spontaneous neuronal activity distinguishes human dorsal and ventral attention systems. Proc Natl Acad Sci U S A. 103:10046–10051.

Fox MD, Snyder AZ, Vincent JL, Corbetta M, Van Essen DC, Raichle ME. 2005. The human brain is intrinsically organized into dynamic, anticorrelated functional networks. Proc Natl Acad Sci U S A. 102:9673–9678.

Foxe JJ, Schroeder CE. 2005. The case for feedforward multisensory convergence during early cortical processing. NeuroReport. 16:419–423.

Frost DO, Moy YP. 1989. Effects of dark rearing on the development of visual callosal connections. Exp Brain Res. 78:203–213.

Gainey MA, Feldman DE. 2017. Multiple shared mechanisms for homeostatic plasticity in rodent somatosensory and visual cortex. Philos Trans R Soc Lond B Biol Sci. 372.

Gamanut R, Kennedy H, Toroczkai Z, Ercsey-Ravasz M, Van Essen DC, Knoblauch K, Burkhalter A. 2018. The Mouse Cortical Connectome, Characterized by an Ultra-Dense Cortical Graph, Maintains Specificity by Distinct Connectivity Profiles. Neuron. 97:698–715 e610.

Gamanut R, Shimaoka D. 2022. Anatomical and functional connectomes underlying hierarchical visual processing in mouse visual system. Brain Struct Funct. 227:1297–1315.

Gao E, DeAngelis GC, Burkhalter A. 2010. Parallel input channels to mouse primary visual cortex. J Neurosci. 30:5912–5926.

Gao M, Maynard KR, Chokshi V, Song L, Jacobs C, Wang H, Tran T, Martinowich K, Lee HK. 2014. Rebound potentiation of inhibition in juvenile visual cortex requires vision-induced BDNF expression. J Neurosci. 34:10770–10779.

Gilissen SR, Arckens L. 2021. Posterior parietal cortex contributions to cross-modal brain plasticity upon sensory loss. Curr Opin Neurobiol. 67:16–25.

Glickfeld LL, Histed MH, Maunsell JH. 2013. Mouse primary visual cortex is used to detect both orientation and contrast changes. J Neurosci. 33:19416–19422.

Glickfeld LL, Olsen SR. 2017. Higher-Order Areas of the Mouse Visual Cortex. Annual Review of Vision Science. 3:251–273.

Godement P, Saillour P, Imbert M. 1979. Thalamic afferents to the visual cortex in congenitally anophthalmic mice. Neurosci Lett. 13:271–278.

Goldreich D, Kanics IM. 2006. Performance of blind and sighted humans on a tactile grating detection task. Percept Psychophys. 68:1363–1371.

Gudi-Mindermann H, Rimmele JM, Nolte G, Bruns P, Engel AK, Roder B. 2018. Working memory training in congenitally blind individuals results in an integration of occipital cortex in functional networks. Behav Brain Res. 348:31–41.

Guimera R, Nunes Amaral LA. 2005. Functional cartography of complex metabolic networks. Nature. 433:895–900.

Hanganu IL, Ben-Ari Y, Khazipov R. 2006. Retinal waves trigger spindle bursts in the neonatal rat visual cortex. J Neurosci. 26:6728–6736.

Harris JA, Mihalas S, Hirokawa KE, Whitesell JD, Choi H, Bernard A, Bohn P, Caldejon S, Casal L, Cho A, Feiner A, Feng D, Gaudreault N, Gerfen CR, Graddis N, Groblewski PA, Henry AM, Ho A, Howard R, Knox JE, Kuan L, Kuang X, Lecoq J, Lesnar P, Li Y, Luviano J, McConoughey S, Mortrud MT, Naeemi M, Ng L, Oh SW, Ouellette B, Shen E, Sorensen SA, Wakeman W, Wang Q, Wang Y, Williford A, Phillips JW, Jones AR, Koch C, Zeng H. 2019. Hierarchical organization of cortical and thalamic connectivity. Nature. 575:195–202.

Harris KD, Mrsic-Flogel TD. 2013. Cortical connectivity and sensory coding. Nature. 503:51–58.

Harris KD, Shepherd GM. 2015. The neocortical circuit: themes and variations. Nat Neurosci. 18:170–181.

Hawellek DJ, Schepers IM, Roeder B, Engel AK, Siegel M, Hipp JF. 2013. Altered intrinsic neuronal interactions in the visual cortex of the blind. J Neurosci. 33:17072–17080.

Heimel JA, Hermans JM, Sommeijer JP, Neuro-Bsik Mouse Phenomics c, Levelt CN. 2008. Genetic control of experience-dependent plasticity in the visual cortex. Genes Brain Behav. 7:915–923.

Heine L, Bahri MA, Cavaliere C, Soddu A, Laureys S, Ptito M, Kupers R. 2015. Prevalence of increases in functional connectivity in visual, somatosensory and language areas in congenital blindness. Front Neuroanat. 9:86.

Hellyer PJ, Jachs B, Clopath C, Leech R. 2016. Local inhibitory plasticity tunes macroscopic brain dynamics and allows the emergence of functional brain networks. NeuroImage. 124:85–95.

Hensch TK. 2005. Critical period mechanisms in developing visual cortex. Curr Top Dev Biol. 69:215–237.

Hindley EL, Nelson AJ, Aggleton JP, Vann SD. 2014. Dysgranular retrosplenial cortex lesions in rats disrupt cross-modal object recognition. Learn Mem. 21:171–179.

Hyvarinen J, Hyvarinen L, Linnankoski I. 1981. Modification of parietal association cortex and functional blindness after binocular deprivation in young monkeys. Exp Brain Res. 42:1–8.

Innocenti GM, Frost DO, Illes J. 1985. Maturation of visual callosal connections in visually deprived kittens: a challenging critical period. J Neurosci. 5:255–267.

Iurilli G, Ghezzi D, Olcese U, Lassi G, Nazzaro C, Tonini R, Tucci V, Benfenati F, Medini P. 2012. Sound-driven synaptic inhibition in primary visual cortex. Neuron. 73:814–828.

Jin L, Lange W, Kempmann A, Maybeck V, Gunther A, Gruteser N, Baumann A, Offenhausser A. 2016. High-efficiency transduction and specific expression of ChR2opt for optogenetic manipulation of primary cortical neurons mediated by recombinant adeno-associated viruses. J Biotechnol. 233:171–180.

Kanjlia S, Lane C, Feigenson L, Bedny M. 2016. Absence of visual experience modifies the neural basis of numerical thinking. Proceedings of the National Academy of Sciences. 113:11172–11177.

Kaschube M, Wolf F, Geisel T, Löwel S. 2002. Genetic Influence on Quantitative Features of Neocortical Architecture. The Journal of Neuroscience. 22:7206–7217.

Keck T, Keller GB, Jacobsen RI, Eysel UT, Bonhoeffer T, Hubener M. 2013. Synaptic scaling and homeostatic plasticity in the mouse visual cortex in vivo. Neuron. 80:327–334.

Klinge C, Eippert F, Roder B, Buchel C. 2010. Corticocortical connections mediate primary visual cortex responses to auditory stimulation in the blind. J Neurosci. 30:12798–12805.

Kraft AW, Mitra A, Bauer AQ, Snyder AZ, Raichle ME, Culver JP, Lee JM. 2017. Visual experience sculpts whole-cortex spontaneous infraslow activity patterns through an Arc-dependent mechanism. Proc Natl Acad Sci U S A. 114:E9952–E9961.

Kriegseis A, Hennighausen E, Rosler F, Roder B. 2006. Reduced EEG alpha activity over parieto-occipital brain areas in congenitally blind adults. Clin Neurophysiol. 117:1560–1573.

Kugler S, Kilic E, Bahr M. 2003. Human synapsin 1 gene promoter confers highly neuron-specific long-term transgene expression from an adenoviral vector in the adult rat brain depending on the transduced area. Gene Ther. 10:337–347.

Kujala T, Alho K, Huotilainen M, Ilmoniemi RJ, Lehtokoski A, Leinonen A, Rinne T, Salonen O, Sinkkonen J, Standertskjold-Nordenstam CG, Naatanen R. 1997. Electrophysiological evidence for cross-modal plasticity in humans with early- and late-onset blindness. Psychophysiology. 34:213–216.

Lambo ME, Turrigiano GG. 2013. Synaptic and Intrinsic Homeostatic Mechanisms Cooperate to Increase L2/3 Pyramidal Neuron Excitability during a Late Phase of Critical Period Plasticity. The Journal of Neuroscience. 33:8810.

Laramée ME, Bronchti G, Boire D. 2014. Primary visual cortex projections to extrastriate cortices in enucleated and anophthalmic mice. Brain Struct Funct. 219:2051–2070.

Laramee ME, Kurotani T, Rockland KS, Bronchti G, Boire D. 2011. Indirect pathway between the primary auditory and visual cortices through layer V pyramidal neurons in V2L in mouse and the effects of bilateral enucleation. Eur J Neurosci. 34:65–78.

Laramée ME, Smolders K, Hu TT, Bronchti G, Boire D, Arckens L. 2016. Congenital Anophthalmia and Binocular Neonatal Enucleation Differently Affect the Proteome of Primary and Secondary Visual Cortices in Mice. PLoS One. 11:e0159320.

Lee H-K, Whitt JL. 2015. Cross-modal synaptic plasticity in adult primary sensory cortices. Current opinion in neurobiology. 35:119–126.

Lee HK. 2022. Metaplasticity framework for cross-modal synaptic plasticity in adults. Front Synaptic Neurosci. 14:1087042.

Lepore N, Voss P, Lepore F, Chou YY, Fortin M, Gougoux F, Lee AD, Brun C, Lassonde M, Madsen SK, Toga AW, Thompson PM. 2010. Brain structure changes visualized in early- and late-onset blind subjects. Neuroimage. 49:134–140.

Lessard N, Pare M, Lepore F, Lassonde M. 1998. Early-blind human subjects localize sound sources better than sighted subjects. Nature. 395:278–280.

Liu Y, Yu C, Liang M, Li J, Tian L, Zhou Y, Qin W, Li K, Jiang T. 2007. Whole brain functional connectivity in the early blind. Brain. 130:2085–2096.

Lohmann G, Margulies DS, Horstmann A, Pleger B, Lepsien J, Goldhahn D, Schloegl H, Stumvoll M, Villringer A, Turner R. 2010. Eigenvector centrality mapping for analyzing connectivity patterns in fMRI data of the human brain. PLoS One. 5:e10232.

Lopez-Bendito G, Anibal-Martinez M, Martini FJ. 2022. Cross-Modal Plasticity in Brains Deprived of Visual Input Before Vision. Annu Rev Neurosci. 45:471–489.

Lubinus C, Orpella J, Keitel A, Gudi-Mindermann H, Engel AK, Roeder B, Rimmele JM. 2021. Data-Driven Classification of Spectral Profiles Reveals Brain Region-Specific Plasticity in Blindness. Cereb Cortex. 31:2505–2522.

Lund RD, Mitchell DE. 1979. The effects of dark-rearing on visual callosal connections of cats. Brain Res. 167:172–175.

Macaluso E. 2006. Multisensory Processing in Sensory-Specific Cortical Areas. The Neuroscientist. 12:327–338.

Macaluso E, Driver J. 2005. Multisensory spatial interactions: a window onto functional integration in the human brain. Trends in Neurosciences. 28:264–271.

Magrou L, Joyce MKP, Froudist-Walsh S, Datta D, Wang XJ, Martinez-Trujillo J, Arnsten AFT. 2024. The meso-connectomes of mouse, marmoset, and macaque: network organization and the emergence of higher cognition. Cereb Cortex. 34.

Makin TR, Krakauer JW. 2023. Against cortical reorganisation. Elife. 12.

Masse IO, Guillemette S, Laramee ME, Bronchti G, Boire D. 2014. Strain differences of the effect of enucleation and anophthalmia on the size and growth of sensory cortices in mice. Brain Res. 1588:113–126.

Matsuzaki Y, Tanaka M, Hakoda S, Masuda T, Miyata R, Konno A, Hirai H. 2019. Neurotropic Properties of AAV-PHP.B Are Shared among Diverse Inbred Strains of Mice. Mol Ther. 27:700–704.

Meister M, Wong RO, Baylor DA, Shatz CJ. 1991. Synchronous bursts of action potentials in ganglion cells of the developing mammalian retina. Science. 252:939–943.

Meng X, Kao JP, Lee HK, Kanold PO. 2015. Visual Deprivation Causes Refinement of Intracortical Circuits in the Auditory Cortex. Cell Rep. 12:955–964.

Meredith MA, Lomber SG. 2017. Species-dependent role of crossmodal connectivity among the primary sensory cortices. Hear Res. 343:83–91.

Michelson NJ, Vanni MP, Murphy TH. 2019. Comparison between transgenic and AAV-PHP.eB-mediated expression of GCaMP6s using in vivo wide-field functional imaging of brain activity. Neurophotonics. 6:025014.

Miller B, Chou L, Finlay BL. 1993. The early development of thalamocortical and corticothalamic projections. J Comp Neurol. 335:16–41.

Missitzi J, Gentner R, Geladas N, Politis P, Karandreas N, Classen J, Klissouras V. 2011. Plasticity in human motor cortex is in part genetically determined. J Physiol. 589:297–306.

Mitra A, Snyder AZ, Hacker CD, Raichle ME. 2014. Lag structure in resting-state fMRI. J Neurophysiol. 111:2374–2391.

Mitra A, Snyder AZ, Tagliazucchi E, Laufs H, Raichle ME. 2015. Propagated infra-slow intrinsic brain activity reorganizes across wake and slow wave sleep. eLife. 4:e10781.

Mooney R, Penn AA, Gallego R, Shatz CJ. 1996. Thalamic relay of spontaneous retinal activity prior to vision. Neuron. 17:863–874.

Morimoto MM, Uchishiba E, Saleem AB. 2021. Organization of feedback projections to mouse primary visual cortex. iScience. 24:102450.

Murphy MC, Nau AC, Fisher C, Kim SG, Schuman JS, Chan KC. 2016. Top-down influence on the visual cortex of the blind during sensory substitution. Neuroimage. 125:932–940.

Noebels JL, Roth WT, Kopell BS. 1978. Cortical slow potentials and the occipital EEG in congenital blindness. J Neurol Sci. 37:51–58.

Noppeney U, Friston KJ, Ashburner J, Frackowiak R, Price CJ. 2005. Early visual deprivation induces structural plasticity in gray and white matter. Curr Biol. 15:R488–490.

Oh SW, Harris JA, Ng L, Winslow B, Cain N, Mihalas S, Wang Q, Lau C, Kuan L, Henry AM, Mortrud MT, Ouellette B, Nguyen TN, Sorensen SA, Slaughterbeck CR, Wakeman W, Li Y, Feng D, Ho A, Nicholas E, Hirokawa KE, Bohn P, Joines KM, Peng H, Hawrylycz MJ, Phillips JW, Hohmann JG, Wohnoutka P, Gerfen CR, Koch C, Bernard A, Dang C, Jones AR, Zeng H. 2014. A mesoscale connectome of the mouse brain. Nature. 508:207–214.

Olavarria J, Bravo H, Ruiz G. 1988. The pattern of callosal connections in posterior neocortex of congenitally anophthalmic rats. Anat Embryol (Berl). 178:155–159.

Olavarria J, Malach R, Van Sluyters RC. 1987. Development of visual callosal connections in neonatally enucleated rats. J Comp Neurol. 260:321–348.

Olavarria J, van Sluyters RC. 1984. Callosal connections of the posterior neocortex in normal-eyed, congenitally anophthalmic, and neonatally enucleated mice. J Comp Neurol. 230:249–268.

Olavarria JF, Li CP. 1995. Effects of neonatal enucleation on the organization of callosal linkages in striate cortex of the rat. J Comp Neurol. 361:138–151.

Ortiz-Terán L, Diez I, Ortiz T, Perez DL, Aragón JI, Costumero V, Pascual-Leone A, El Fakhri G, Sepulcre J. 2017. Brain circuit-gene expression relationships and neuroplasticity of multisensory cortices in blind children. Proc Natl Acad Sci U S A. 114:6830–6835.

Oude Lohuis MN, Marchesi P, Olcese U, Pennartz CMA. 2024. Triple dissociation of visual, auditory and motor processing in mouse primary visual cortex. Nature Neuroscience. 27:758–771.

Petrus E, Isaiah A, Jones Adam P, Li D, Wang H, Lee H-K, Kanold Patrick O. 2014. Crossmodal Induction of Thalamocortical Potentiation Leads to Enhanced Information Processing in the Auditory Cortex. Neuron. 81:664–673.

Petrus E, Isaiah A, Jones AP, Li D, Wang H, Lee HK, Kanold PO. 2014. Crossmodal induction of thalamocortical potentiation leads to enhanced information processing in the auditory cortex. Neuron. 81:664–673.

Petrus E, Rodriguez G, Patterson R, Connor B, Kanold PO, Lee HK. 2015. Vision loss shifts the balance of feedforward and intracortical circuits in opposite directions in mouse primary auditory and visual cortices. J Neurosci. 35:8790–8801.

Piché M, Robert S, Miceli D, Bronchti G. 2004. Environmental enrichment enhances auditory takeover of the occipital cortex in anophthalmic mice. Eur J Neurosci. 20:3463–3472.

Poirier C, Collignon O, Scheiber C, Renier L, Vanlierde A, Tranduy D, Veraart C, De Volder AG. 2006. Auditory motion perception activates visual motion areas in early blind subjects. Neuroimage. 31:279–285.

Power JD, Cohen AL, Nelson SM, Wig GS, Barnes KA, Church JA, Vogel AC, Laumann TO, Miezin FM, Schlaggar BL, Petersen SE. 2011. Functional network organization of the human brain. Neuron. 72:665–678.

Ptito M, Matteau I, Zhi Wang A, Paulson OB, Siebner HR, Kupers R. 2012. Crossmodal recruitment of the ventral visual stream in congenital blindness. Neural Plast. 2012:304045.

Ptito M, Schneider FC, Paulson OB, Kupers R. 2008. Alterations of the visual pathways in congenital blindness. Exp Brain Res. 187:41–49.

Qin W, Liu Y, Jiang T, Yu C. 2013. The development of visual areas depends differently on visual experience. PLoS One. 8:e53784.

Renier L, Cuevas I, Grandin CB, Dricot L, Plaza P, Lerens E, Rombaux P, De Volder AG. 2013. Right occipital cortex activation correlates with superior odor processing performance in the early blind. PLoS One. 8:e71907.

Rhoades RW, Mooney RD, Fish SE. 1984. A comparison of visual callosal organization in normal, bilaterally enucleated and congenitally anophthalmic mice. Exp Brain Res. 56:92–105.

Rhoades RW, Mooney RD, Fish SE. 1985. Subcortical projections of area 17 in the anophthalmic mouse. Brain Res. 349:171–181.

Ricciardi E, Bonino D, Pellegrini S, Pietrini P. 2014. Mind the blind brain to understand the sighted one! Is there a supramodal cortical functional architecture? Neurosci Biobehav Rev. 41:64–77.

Ricciardi E, Tozzi L, Leo A, Pietrini P. 2014. Modality dependent cross-modal functional reorganization following congenital visual deprivation within occipital areas: a meta-analysis of tactile and auditory studies. Multisens Res. 27:247–262.

Rimmele JM, Gudi-Mindermann H, Nolte G, Röder B, Engel AK. 2019. Working memory training integrates visual cortex into beta-band networks in congenitally blind individuals. Neuroimage. 194:259–271.

Rocha RP, Koçillari L, Suweis S, Corbetta M, Maritan A. 2018. Homeostatic plasticity and emergence of functional networks in a whole-brain model at criticality. Scientific Reports. 8:15682.

Roder B, Rosler F, Hennighausen E, Nacker F. 1996. Event-related potentials during auditory and somatosensory discrimination in sighted and blind human subjects. Brain Res Cogn Brain Res. 4:77–93.

Roder B, Teder-Salejarvi W, Sterr A, Rosler F, Hillyard SA, Neville HJ. 1999. Improved auditory spatial tuning in blind humans. Nature. 400:162–166.

Rubinov M, Sporns O. 2010. Complex network measures of brain connectivity: uses and interpretations. Neuroimage. 52:1059–1069.

Saccone EJ, Tian M, Bedny M. 2024. Developing cortex is functionally pluripotent: Evidence from blindness. Dev Cogn Neurosci. 66:101360.

Sadato N, Pascual-Leone A, Grafman J, Deiber MP, Ibanez V, Hallett M. 1998. Neural networks for Braille reading by the blind. Brain. 121 ( Pt 7):1213–1229.

Sadato N, Pascual-Leone A, Grafman J, Ibanez V, Deiber MP, Dold G, Hallett M. 1996. Activation of the primary visual cortex by Braille reading in blind subjects. Nature. 380:526–528.

Saleem AB. 2020. Two stream hypothesis of visual processing for navigation in mouse. Curr Opin Neurobiol. 64:70–78.

Schepers IM, Hipp JF, Schneider TR, Roder B, Engel AK. 2012. Functionally specific oscillatory activity correlates between visual and auditory cortex in the blind. Brain. 135:922–934.

Schmidt R, LaFleur KJ, de Reus MA, van den Berg LH, van den Heuvel MP. 2015. Kuramoto model simulation of neural hubs and dynamic synchrony in the human cerebral connectome. BMC Neurosci. 16:54.

Schubert JT, Buchholz VN, Focker J, Engel AK, Roder B, Heed T. 2015. Oscillatory activity reflects differential use of spatial reference frames by sighted and blind individuals in tactile attention. Neuroimage. 117:417–428.

Sforazzini F, Schwarz AJ, Galbusera A, Bifone A, Gozzi A. 2014. Distributed BOLD and CBV-weighted resting-state networks in the mouse brain. Neuroimage. 87:403–415.

Shu N, Liu Y, Li J, Li Y, Yu C, Jiang T. 2009. Altered anatomical network in early blindness revealed by diffusion tensor tractography. PLoS One. 4:e7228.

Smolders K, Vreysen S, Laramée M-E, Cuyvers A, Hu T-T, Van Brussel L, Eysel UT, Nys J, Arckens L. 2016. Retinal lesions induce fast intrinsic cortical plasticity in adult mouse visual system. European Journal of Neuroscience. 44:2165–2175.

Striem-Amit E, Ovadia-Caro S, Caramazza A, Margulies DS, Villringer A, Amedi A. 2015. Functional connectivity of visual cortex in the blind follows retinotopic organization principles. Brain. 138:1679–1695.

Tian M, Xiao X, Hu H, Cusack R, Bedny M. 2023. Visual experience shapes functional connectivity between occipital and non-visual networks. bioRxiv.

Tomaiuolo F, Campana S, Collins DL, Fonov VS, Ricciardi E, Sartori G, Pietrini P, Kupers R, Ptito M. 2014. Morphometric changes of the corpus callosum in congenital blindness. PLoS One. 9:e107871.

Touj S, Tokunaga R, Al Ain S, Bronchti G, Piche M. 2019. Pain Hypersensitivity is Associated with Increased Amygdala Volume and c-Fos Immunoreactivity in Anophthalmic Mice. Neuroscience. 418:37–49.

Tucker P, Laemle L, Munson A, Kanekar S, Oliver ER, Brown N, Schlecht H, Vetter M, Glaser T. 2001. The eyeless mouse mutation (ey1) removes an alternative start codon from the Rx/rax homeobox gene. Genesis. 31:43–53.

Turley JA, Zalewska K, Nilsson M, Walker FR, Johnson SJ. 2017. An analysis of signal processing algorithm performance for cortical intrinsic optical signal imaging and strategies for algorithm selection. Scientific Reports. 7:7198.

Valley MT, Moore MG, Zhuang J, Mesa N, Castelli D, Sullivan D, Reimers M, Waters J. 2020. Separation of hemodynamic signals from GCaMP fluorescence measured with wide-field imaging. Journal of Neurophysiology. 123:356–366.

van den Heuvel MP, Sporns O. 2011. Rich-club organization of the human connectome. J Neurosci. 31:15775–15786.

van der Heijden K, Formisano E, Valente G, Zhan M, Kupers R, de Gelder B. 2019. Reorganization of Sound Location Processing in the Auditory Cortex of Blind Humans. Cerebral Cortex. 30:1103–1116.

Vanhatalo S, Palva JM, Holmes MD, Miller JW, Voipio J, Kaila K. 2004. Infraslow oscillations modulate excitability and interictal epileptic activity in the human cortex during sleep. Proc Natl Acad Sci U S A. 101:5053–5057.

Wang D, Qin W, Liu Y, Zhang Y, Jiang T, Yu C. 2014. Altered resting-state network connectivity in congenital blind. Hum Brain Mapp. 35:2573–2581.

Wang Q, Sporns O, Burkhalter A. 2012. Network Analysis of Corticocortical Connections Reveals Ventral and Dorsal Processing Streams in Mouse Visual Cortex. The Journal of Neuroscience. 32:4386–4399.

Watkins KE, Cowey A, Alexander I, Filippini N, Kennedy JM, Smith SM, Ragge N, Bridge H. 2012. Language networks in anophthalmia: maintained hierarchy of processing in ‘visual’ cortex. Brain. 135:1566–1577.

Weeks R, Horwitz B, Aziz-Sultan A, Tian B, Wessinger CM, Cohen LG, Hallett M, Rauschecker JP. 2000. A positron emission tomographic study of auditory localization in the congenitally blind. J Neurosci. 20:2664–2672.

Weliky M, Katz LC. 1999. Correlational structure of spontaneous neuronal activity in the developing lateral geniculate nucleus in vivo. Science. 285:599–604.

Wittenberg G, Werhahn K, Wassermann E, Herscovitch P, Cohen L. 2004. Functional connectivity between somatosensory and visual cortex in ear ly blind humans. European Journal of Neuroscience.

Wong RO, Meister M, Shatz CJ. 1993. Transient period of correlated bursting activity during development of the mammalian retina. Neuron. 11:923–938.

Yu C, Liu Y, Li J, Zhou Y, Wang K, Tian L, Qin W, Jiang T, Li K. 2008. Altered functional connectivity of primary visual cortex in early blindness. Hum Brain Mapp. 29:533–543.

Zingg B, Hintiryan H, Gou L, Song MY, Bay M, Bienkowski MS, Foster NN, Yamashita S, Bowman I, Toga AW, Dong H-W. 2014. Neural networks of the mouse neocortex. Cell. 156:1096–1111.

