## Supplementary Material for "Cortical Functional Connectivity in Mouse Models of Early Blindness: Enucleation vs. Anophthalmia"

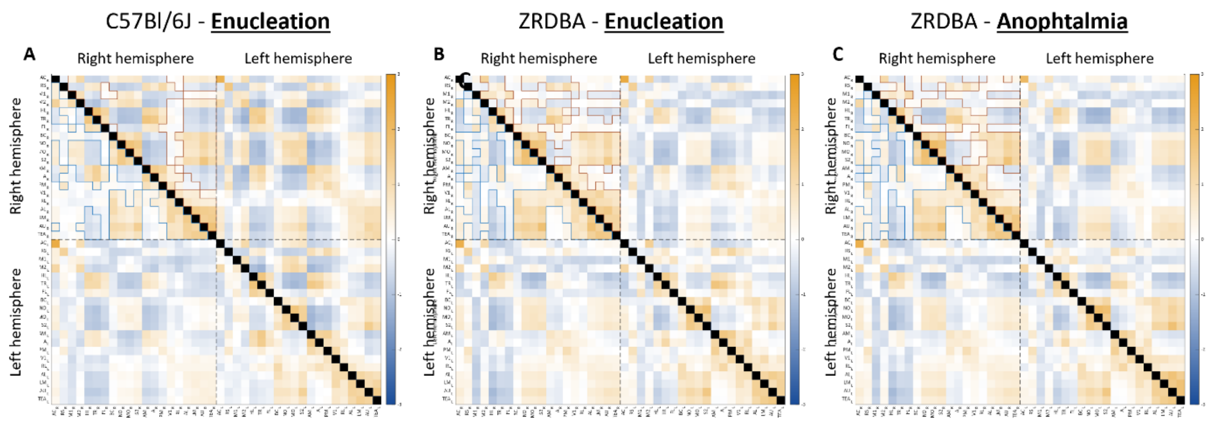


Supplementary Figure 1 - Functional connectivity matrix between Sighted and Blind (A) C57Bl/6J and (B–C) ZRDBA mice.

Average Fisher z-transformed correlation matrix illustrating pairwise functional connectivity across 40 bilateral cortical regions of interest (ROIs). The lower triangle displays data from the sighted group; the upper triangle corresponds to the (A–B) enucleated and (C) anophthalmic mice. Autologous connections along the main diagonal are masked in black. ROIs are grouped by hemisphere (right: ROIs 1–20; left: ROIs 21–40).

The matrix is visually divided into four quadrants delimited by the dashed lines: quadrants I (top-left) and IV (bottom right) correspond to intrahemispheric connectivity within the right and left hemispheres, respectively, while quadrant II (top right) and III (bottom-left) represent interhemispheric connectivity between homologous or heterologous ROIs across hemispheres.

Color contours overlay the right hemisphere to visualize absolute functional connectivity above 0.3 (absolute threshold), used to construct the adjacency matrices for each group's connectome. Blue outlines correspond to the adjacency matrix of the sighted group, and brown outlines to those of the blind groups.

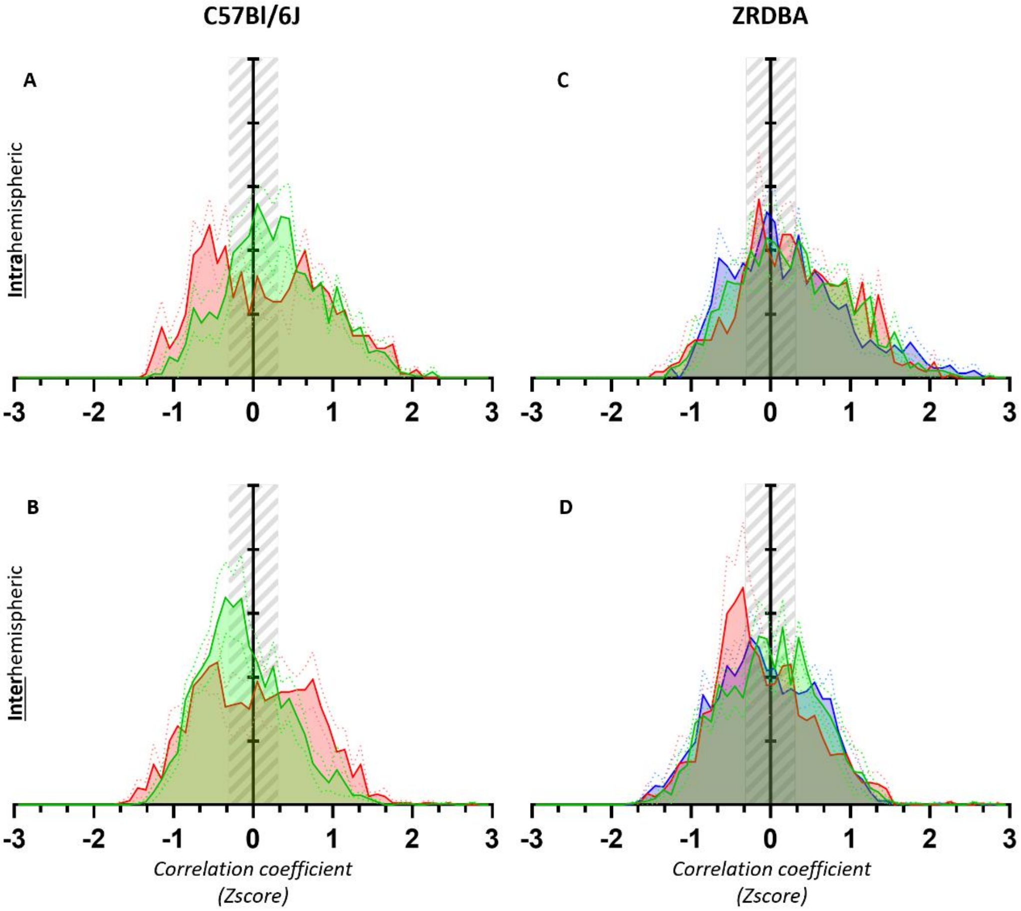


**Supplementary Figure 2 - Frequency distribution of Normalized Pearson's Correlation Coefficients for Infraslow Oscillations**

Pearson's correlation coefficient distributions for infraslow oscillations (<0.1 Hz) during resting-state widefield calcium imaging between intrahemispheric (A-C) and interhemispheric (B-D) cortical areas. Solid lines indicate group means, dashed lines show SEM, and the light grey-hatched rectangle highlights the threshold (zscore Pearson's correlation ≥ 0.3) applied for connectomic analysis

Intrahemispheric : Comparisons are shown between sighted (green) and enucleated (red) C57Bl/6J mice (**A**) and between sighted (green), enucleated (red), and anophthalmic (blue) ZRDBA mice (**C**).

Interhemispheric : Results are presented for sighted and enucleated C57Bl/6J mice (**B**) and for sighted, enucleated, and anophthalmic ZRDBA mice (**D**).

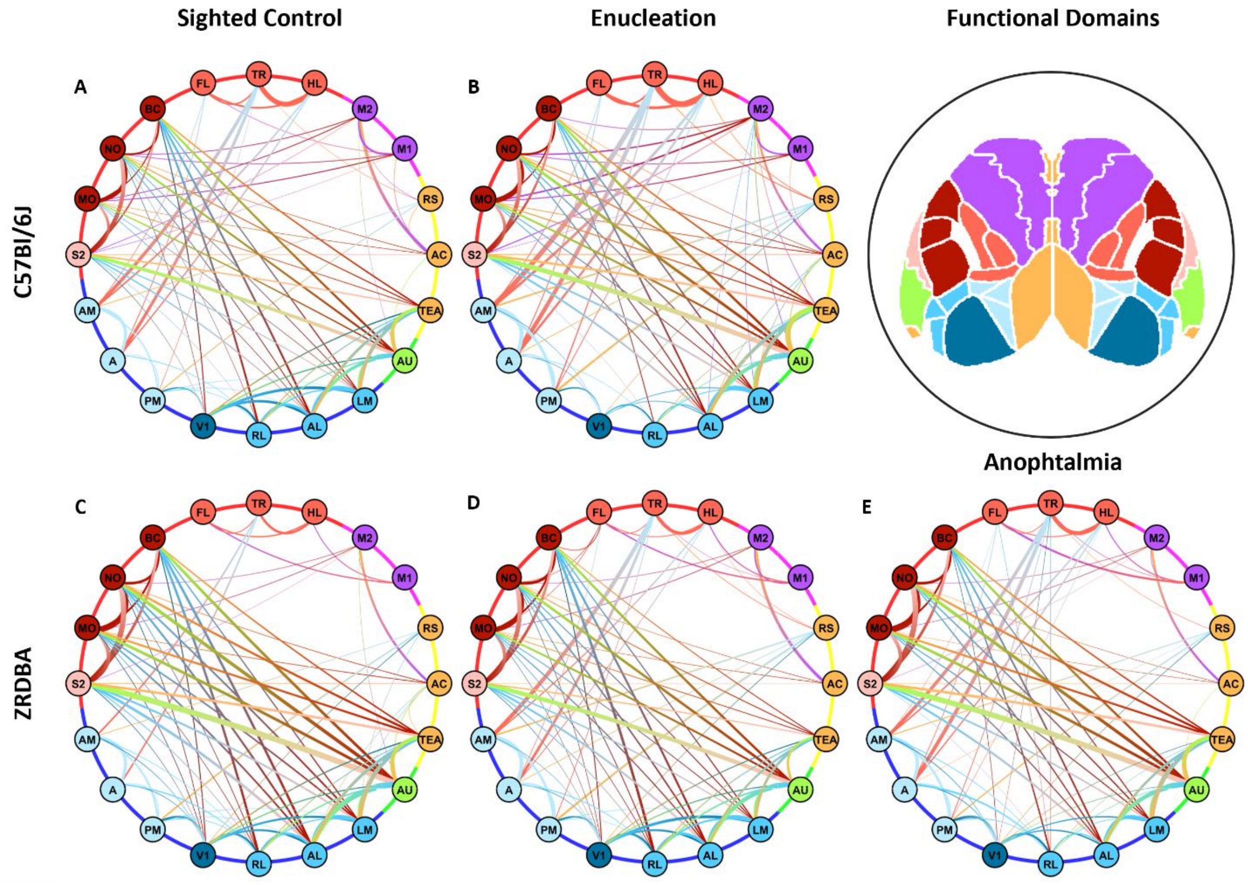


**Supplementary Figure 3 - Circular connectivity graphs illustrating cortico-cortical functional interactions** in five experimental groups: **(A)** *Sighted*, and **(B)** *Enucleated C57Bl/6J mice*, and **(C)** *Sighted*, **(D)** *Enucleated* and **(E)** *Anophthalmic ZRDBA mice*. Each node represents a distinct cortical region of interest (ROI), color-coded by functional modality: associative (nectarine), motor (crocus), lemniscal (firebrick), secondary somatosensory (marshmallow), trigeminal (paprika), medial visual (meltwater), primary visual V1 (denim), lateral visual (ocean), and auditory (meadow). Edges represent inter-regional correlations derived from z-score normalized functional connectivity matrices. Only suprathreshold connections (r>0.3) are shown to exclude weak associations. Edge thickness is scaled proportionally to the strength of the connections.
